# The bacterial iron sensor IdeR recognizes its DNA targets by indirect readout

**DOI:** 10.1101/2021.02.12.430932

**Authors:** Francisco Javier Marcos-Torres, Dirk Maurer, Julia J. Griese

## Abstract

IdeR is the main transcriptional regulator controlling iron homeostasis genes in Actinobacteria, including species from the *Corynebacterium, Mycobacterium*, and *Streptomyces* genera, as well as the erythromycin-producing bacterium *Saccharopolyspora erythraea.* Despite being a well-studied transcription factor since the identification of the Diphtheria toxin repressor DtxR three decades ago, the details of how IdeR proteins recognize their highly conserved 19-bp DNA target remain to be elucidated. The results of our structural and mutational studies support a model wherein IdeR uses an indirect readout mechanism, identifying its targets via the sequence-specific DNA backbone structure rather than through direct contacts with the DNA bases. Furthermore, we show that IdeR efficiently recognizes a shorter palindromic sequence corresponding to a half binding site as compared to the full 19-bp target previously reported, expanding the number of potential target genes controlled by IdeR proteins.

## INTRODUCTION

The study of transcriptional regulation in bacteria is critical to our understanding of how microorganisms sense stimuli and how the expression of relevant genes is regulated to adapt to new environmental conditions. *Saccharopolyspora erythraea* is a soil-dwelling actinobacterium best known for producing the macrolide antibiotic erythromycin (Oliynyk *et al*, 2007; Wu *et al*, 2015; Kim & Goodfellow, 2015). Due to the size of its genome and its multicellular behavior, *S. erythraea* is an excellent bacterial system to study genetic regulation in complex organisms. In the highly variable environment of the soil, bacteria have developed a wide range of sensor systems and signal transduction mechanisms to genetically respond to the stimuli detected by those sensors, for instance, if an essential micro- or macro-nutrient is limiting or if a toxic compound is present.

Bacterial signaling systems have traditionally been classified in four groups, known as the four pillars of signal transduction mechanisms, based on the distribution of their sensory input domains and their DNA-binding effector domains. The simplest signal transduction mechanism is exemplified by the group of one-component systems (OCS), where the sensor and the effector DNA-binding domains are part of the same protein (Ulrich *et al*, 2005; Staroń *et al*, 2009; Muñoz-Dorado *et al*, 2012; Marcos-Torres *et al*, 2016; Pinto *et al*, 2019). With a size of 8.2 Mb, the *S. erythraea* genome encodes 675 DNA-binding proteins predicted to be involved in signal transduction, 652 of which are expected to function as OCS (Gumerov *et al*, 2020). Compared to other less complex bacterial organisms such as *Escherichia coli*, with approximately 250 DNA binding proteins classified as OCS (Gumerov *et al*, 2020), *S. erythraea* displays a rich assortment of regulatory proteins. Ideally, to avoid cross-talk between all the transcriptional regulators, every DNA-binding protein should specifically recognize a particular target sequence, but in reality, most of these transcriptional regulators recognize different target DNA sequences that resemble a consensus sequence without perfectly matching it. In a bacterium which possesses so many predicted transcriptional regulators, it is of great interest to understand where the boundaries of this pattern recognition flexibility lie and how it can be manipulated to improve or redirect gene regulation. *S. erythraea* in particular is of great biotechnological interest because it produces erythromycin, and much effort has been made to improve production of this secondary metabolite, yet the complex regulation of this process is not completely understood despite decades of research (Wu *et al*, 2019; Pan *et al*, 2019; Liu *et al*, 2019; Xu *et al*, 2019; Li *et al*, 2020a, 2020b).

One of the most interesting regulatory processes in bacteria is the one controlling iron homeostasis. As life evolved on Earth, iron became an essential element to almost all organisms because in the primitive oxygen-free environment this transition metal was abundant and soluble in its ferrous (Fe^2+^) form. Iron was incorporated into a variety of enzyme cofactors, since it can be used to transfer electrons, act as a Lewis acid, or catalyze redox reactions. As the atmosphere became oxygenated, two major inconveniences arose to which iron-dependent organisms had to adapt. First, iron bioavailability was drastically reduced in all oxygenic environments, as its oxidized ferric (Fe^3+^) form is poorly soluble. Second, iron became highly toxic due to the generation of reactive oxygen species (ROS) through Fenton reactions. Adaptation to these new conditions led to the development of iron uptake and storage mechanisms, including the production of siderophores to complex ferric ions, and to a tight regulation of such mechanisms to avoid the toxicity derived from an excess of iron (Imlay, 2013; Moraleda - Muñoz *et al*, 2019; Guth-Metzler *et al*, 2020). In Gram-negative and Gram-positive bacteria with low GC content, this regulation is usually accomplished by the ferric uptake regulator Fur, whereas in Gram-positive bacteria with high GC content, as well as in archaea, iron homeostasis is frequently controlled by its functional homologue IdeR (iron-dependent regulator) (Hantke, 2001; Leyn & Rodionov, 2015).

IdeR is an OCS from the DtxR (Diphtheria toxin repressor) family of transcriptional regulators which typically consist of three domains: an N-terminal DNA-binding helix-turn-helix (HTH) motif, followed by a dimerization interface that contains most of the metal-binding residues, and a C-terminal SH3-like domain (Qiu *et al*, 1995; Wisedchaisri *et al*, 2004; Granger *et al*, 2013). The main function of IdeR is to repress the expression of iron uptake genes when the intracellular iron levels are sufficient to avoid its toxicity. When intracellular iron levels are low, metal-free IdeR remains inactive, and all iron uptake genes are expressed. When iron levels are sufficient for iron ions to occupy the IdeR metal-binding sites, the regulator is activated and recognizes a highly conserved 19-bp sequence located in the promoter region of its target genes. The binding of IdeR to the promoter generally blocks the transcription of the regulated genes, as is the case for iron uptake genes, thereby preventing the iron concentration from reaching toxic levels. On the other hand, it has also been reported that IdeR can activate the transcription of some genes, such as iron storage genes, in response to high iron concentrations (Gold *et al*, 2001; Pandey & Rodriguez, 2014; Deng & Zhang, 2015; Kurthkoti *et al*, 2015; Cheng *et al*, 2018; Zondervan *et al*, 2018).

The structural and functional details of the DNA-binding mechanism of this regulator are still not fully understood. By screening the ability of IdeR from *S. erythraea* (*Se*IdeR) to bind to variations of a DNA target, and analyzing the structural details of those interactions, we provide an in-depth description of the specificity of this transcriptional regulator and the thresholds of its tolerance for recognizing a particular DNA pattern, unveiling which regulator residues are involved in this process and which DNA bases provide the specific fingerprint being recognized. We show that IdeR recognizes half binding sites, expanding the already vast repertoire of putative binding sites for this type of regulator. We also provide evidence that IdeR uses an indirect readout mechanism to recognize its DNA targets, identifying them by their sequence-dependent backbone structure rather than through contacts with the DNA bases themselves. The similarities of the HTH motif with other IdeR proteins implies that other members of this family of bacterial transcriptional regulators also use an indirect readout mechanism to find their targets.

## RESULTS

### IdeR likely controls the expression of at least 23 gene clusters in *S. erythraea*

The DNA-binding HTH motif of *Se*IdeR shares an amino acid identity of around 92%, 96% and 100% with those of IdeR from *Mycobacterium tuberculosis* and *Streptomyces avermitilis*, and DtxR from *Corynebacterium diphtheriae*, respectively. Considering that these proteins recognize a similar 19-bp DNA target in their respective hosts (Wisedchaisri *et al*, 2004; Cheng *et al*, 2018; Lee & Holmes, 2000), we used the 19-bp consensus sequence to screen the genome of *S. erythraea* for putative IdeR targets. This search provided more than 70 sequences which were manually curated to obtain 37 reliable putative binding sites that could be matched to 23 gene clusters (Table EV1).

Most of the identified gene clusters are involved in the uptake and storage of iron, with a high presence of genes coding for siderophore production or transport, iron ABC transport systems and bacterioferritins, as well as an EfeO-like ferric iron uptake transporter. Additionally, IdeR appears to regulate several gene clusters encoding proteins that use iron as a cofactor. Among these we find ferredoxins, a L-lactate dehydrogenase, and the Nuo NADH dehydrogenase complex of the respiratory chain (Table EV1).

To confirm some of the putative binding sites listed in Table EV1, we analyzed the binding of IdeR to two of the binding sites found in cluster 10. This cluster is predicted to be involved in the production and transport of a siderophore to capture environmental iron. The first two putative IdeR binding sites in the cluster (C10S1 and C10S2) can be found in the intergenic region between genes SACE_2689 and SACE_2690. The orientation of both genes suggests the presence of a divergent promoter in this intergenic region, with C10S1 being closer to the start codon of gene SACE_2689, and C10S2 to the start codon of SACE_2690. To assess IdeR binding to its target sequences in different metalation conditions, we performed an electrophoretic mobility shift assay (EMSA) using 30 nM of fluorescein-labelled C10S1 DNA in the presence of 25 times excess of IdeR and different metal concentrations. As can be seen in Fig 1A, IdeR binds to C10S1 in the presence of Co^2+^, Mn^2+^, and Fe^2+^ starting at concentrations of 15 μM. However, the activation by Mn^2+^ does not appear to be as efficient as by Co^2+^ or Fe^2+^, which activate IdeR at similar concentrations, as previously reported for *M. tuberculosis* IdeR and *C. diphtheriae* DtxR (Schmitt *et al*, 1995; Schmitt & Holmes, 1993). Due to the similarities between both metals (Bertini & Luchinat, 1984), and the constraints of working with Fe^2+^, we used Co^2+^ to activate IdeR in all of the subsequent DNA-binding experiments.

**Figure 1.**
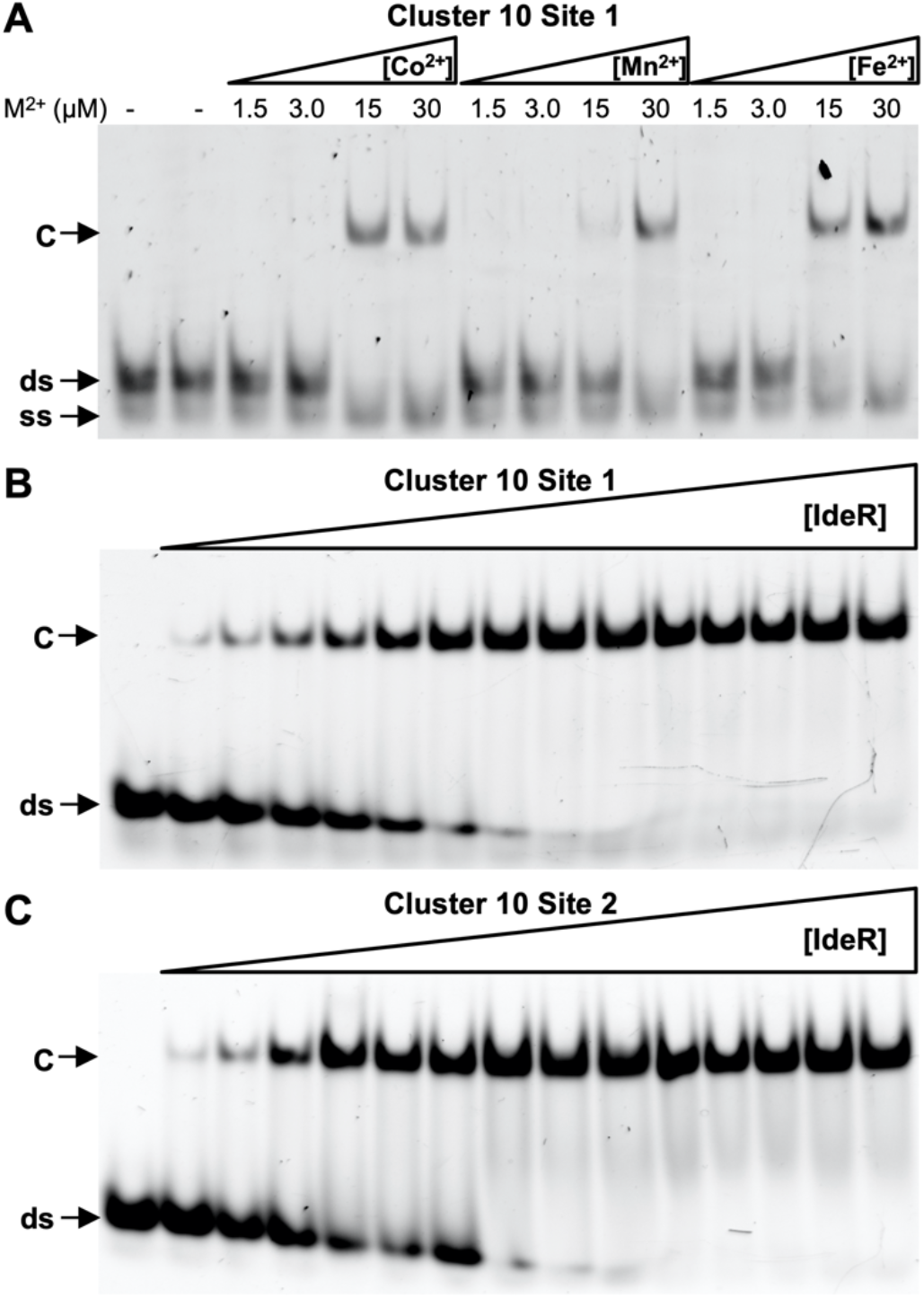
EMSA analysis of IdeR binding to native *S. erythraea* DNA-binding sites. A Binding of IdeR at 750 nM (dimer concentration) to site 1 of cluster 10 in the presence of increasing concentrations of Co^2+^, Mn^2+^ or Fe^2+^. B Binding of IdeR, in increasing concentrations (15 nM - 2.25 μM dimer), to site 1 of cluster 10 in the presence of 30 μM Co^2+^. B Binding of IdeR, in increasing concentrations (15 nM - 2.25 μM dimer), to site 2 of cluster 10 in the presence of 30 μM Co^2+^. IdeR was added to 30 nM fluorescence-labeled double-stranded DNA probe in the presence of competitor DNA. Protein-DNA complexes were resolved on a 4% Tris-acetate polyacrylamide gel. The left-most lane is a control reaction without protein. ds, unbound double-stranded DNA probe; ss, nonhybridized single-stranded DNA; C, protein-dsDNA complex.

The affinity of Co^2+^-activated IdeR for the C10S1 and C10S2 DNA targets was assessed by EMSA using increasing concentrations of IdeR. IdeR binds specifically to both DNA sequences, with similar dissociation constants (*K_D_*), which are estimated to be around 116 nM and 94 nM for the C10S1 and C10S2 targets, respectively (Fig 1B and C, Table 1). These *K_D_* values are in good agreement with those previously reported for IdeR from *M. tuberculosis* (Chou *et al*, 2004; Wisedchaisri *et al*, 2007).

**Table 1.**
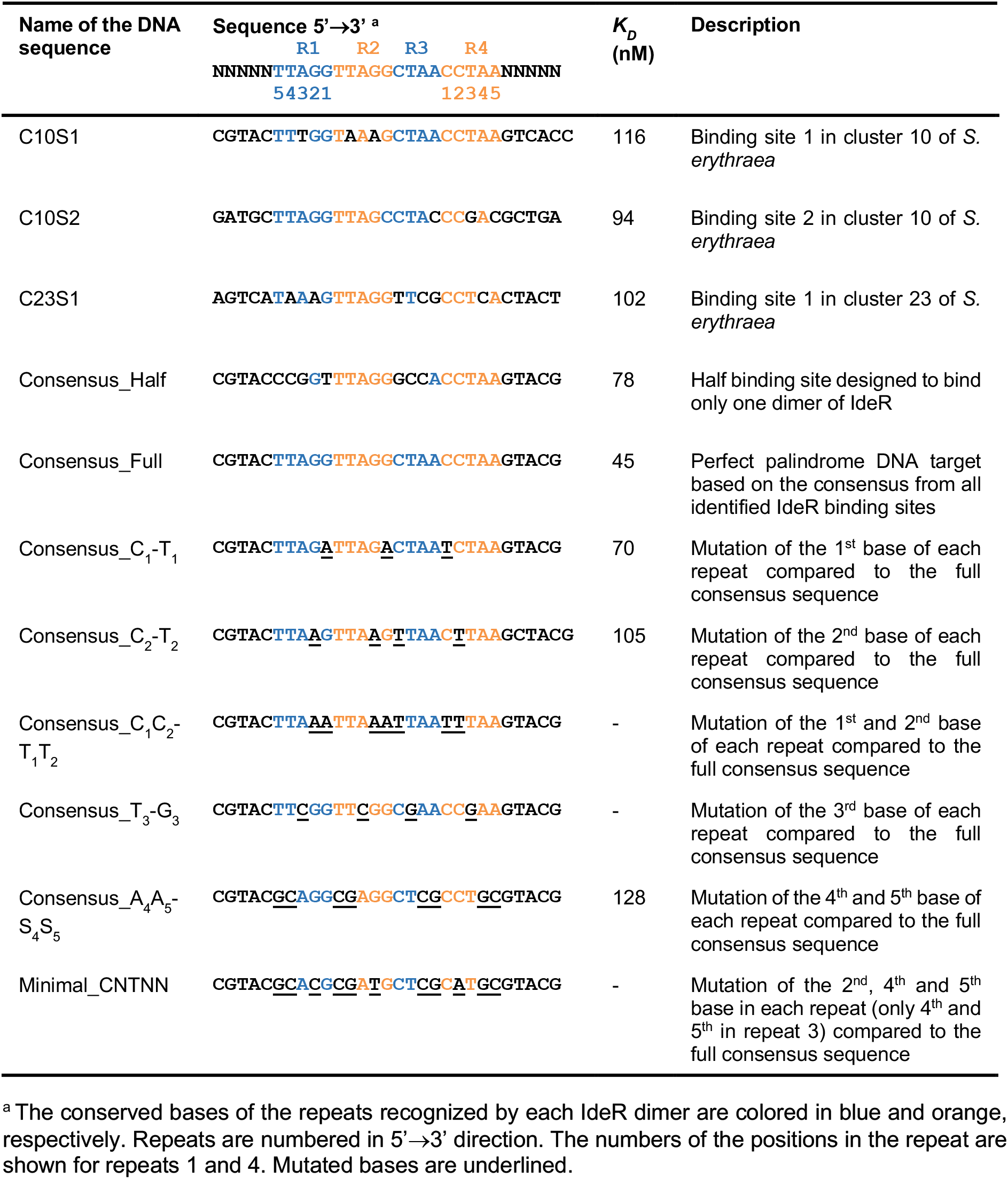
DNA sequences used for EMSA analysis and estimated dissociation constant (*K_D_*) for each binding reaction with IdeR^WT^.

### Two IdeR dimers bind to the palindromic recognition sequence

We determined the crystal structure of Co^2+^-activated IdeR at 2.0 Å resolution (Table 2, Fig EV1A, see also Fig 2A). The protein forms a dimer. Each subunit consists of the three domains typical of an IdeR protein, an N-terminal domain containing the DNA-binding HTH motif, a dimerization domain, and a C-terminal SH3-like domain. Two metal-binding sites are formed by residues from all three domains, coordinating two Co^2+^ ions in octahedral geometry (Fig EV2A, see also Fig 2B). The overall structure of *Se*IdeR is highly similar to the structures of IdeR from *M. tuberculosis* (Wisedchaisri *et al*, 2004) and DtxR from *C. diphtheriae* (Pohl *et al*, 1999) (Appendix Table S1).

**Figure 2.**
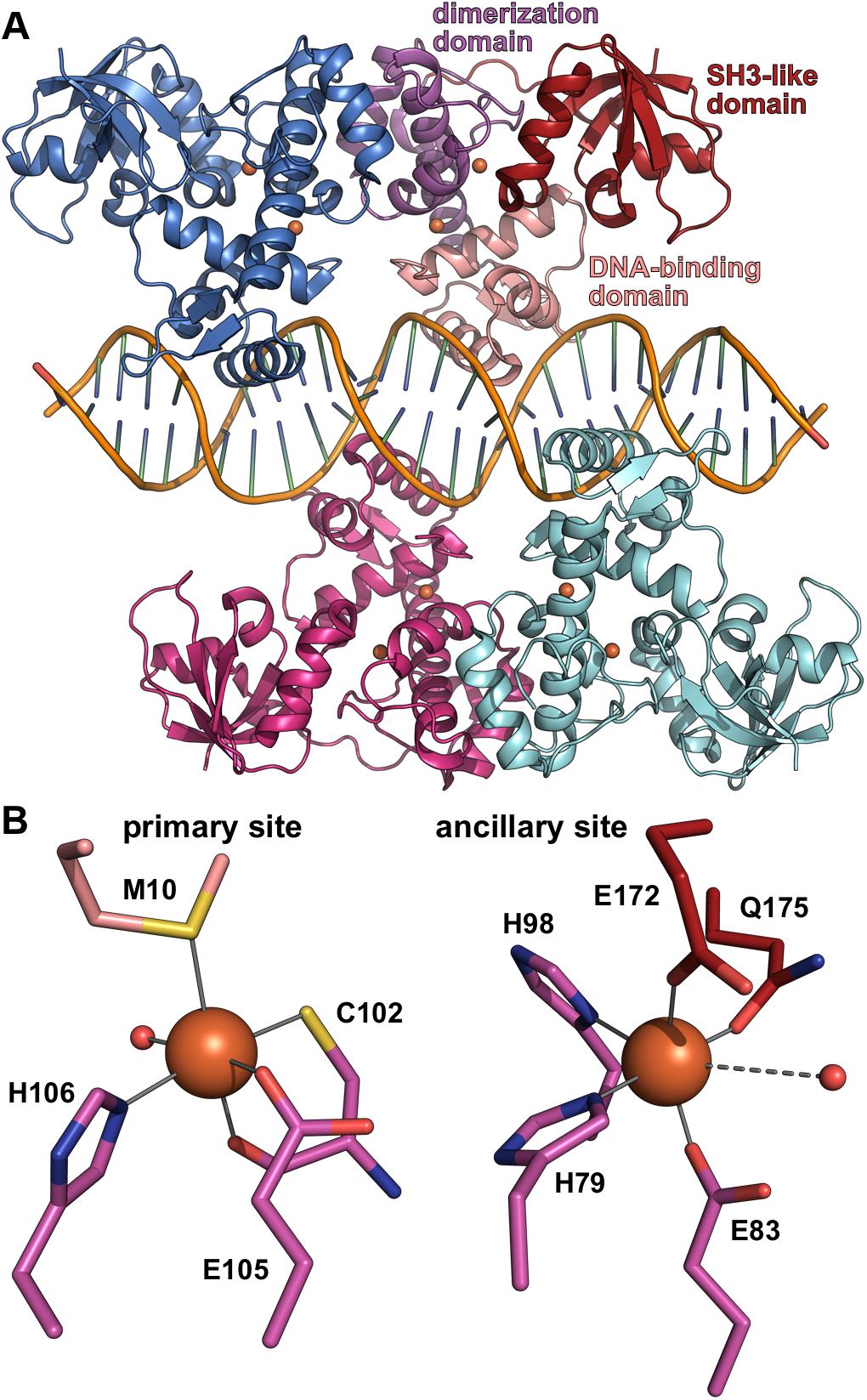
Crystal structure of Fe^2+^-activated IdeR in complex with the consensus DNA-binding sequence. A Overall structure of the complex, with IdeR subunit B colored by domain and the other IdeR subunits colored by subunit. The Fe^2+^ ions are shown as orange spheres. Two IdeR dimers bind to opposite faces of the DNA double helix. B The metal-binding sites in IdeR subunit B, depicted using the same coloring scheme as in A, with water ligands shown as small red spheres. Metal-ligand bonds are indicated by grey lines, the dashed line between the ancillary site metal ion and water ligand indicating a long, weak bond.

**Table 2.** Crystallographic data and refinement statistics.

| Structure | Co<sup>2+</sup>-IdeR<sup>WT</sup> | Co<sup>2+</sup>-IdeR<sup>WT</sup> +<br>consensus<br>DNA | Fe<sup>2+</sup>-IdeR<sup>WT</sup> +<br>consensus<br>DNA | Co<sup>2+</sup>-IdeR<sup>WT</sup> +<br>C10S1 DNA | Co<sup>2+</sup>-IdeR<sup>Q43</sup> +<br>consensus<br>DNA | Co<sup>2+</sup>-IdeR<sup>P39</sup> +<br>consensus<br>DNA |
| --- | --- | --- | --- | --- | --- | --- |
| PDB ID | 7B1V | 7B1Y | 7B20 | 7B23 | 7B25 | 7B24 |
| **Data collection and processing** |  |  |  |  |  |  |
| Wavelength | 0.980 | 0.976 | 0.976 | 0.918 | 0.976 | 0.976 |
| Resolution (Å) | 61.10-2.04<br>(2.22-2.04) | 78.94-2.12<br>(2.32-2.12) | 54.29-2.18<br>(2.44-2.18) | 94.63-2.15<br>(2.41-2.15) | 93.16-2.34<br>(2.63-2.34) | 54.93-2.05<br>(2.23-2.05) |
| Resolution limits along<br>axes (a*, b*, c*) | 2.58, 2.04,<br>2.21 | 2.12, 2.18,<br>3.06 | 2.41, 2.18,<br>2.95 | 2.15, 2.48,<br>2.72 | 2.32, 2.81,<br>3.34 | 2.05, 2.13,<br>2.65 |
| Space group | P1 | C2 | C2 | C2 | C2 | C2 |
| Cell dimensions |  |  |  |  |  |  |
| a, b, c (Å) | 43.88, 45.56,<br>66.28 | 195.04,<br>112.90, 88.51 | 194.12,<br>112.66, 88.55 | 194.25,<br>113.13, 89.29 | 192.66,<br>110.86, 86.75 | 196.40,<br>113.84, 89.80 |
| α, β, γ (°) | 109.12, 93.20,<br>113.98 | 90.00, 117.07,<br>90.00 | 90.00, 117.28,<br>90.00 | 90.00, 117.25,<br>90.00 | 90.00,<br>116.849, 90.00 | 90.00, 117.25,<br>90.00 |
| No. unique reflections | 17216 (862) | 60545 (3027) | 54157 (2708) | 59657 (2984) | 36928 (1847) | 76685 (3835) |
| Multiplicity | 3.3 (2.4) | 6.9 (6.7) | 6.9 (6.7) | 6.7 (6.7) | 6.8 (6.5) | 7.0 (6.9) |
| R<sub>merge</sub> | 0.058 (0.579) | 0.092 (1.203) | 0.093 (1.136) | 0.083 (0.984) | 0.130 (1.218) | 0.082 (1.305) |
| R<sub>meas</sub> | 0.069 (0.748) | 0.099 (1.304) | 0.100 (1.229) | 0.090 (1.066) | 0.141 (1.323) | 0.088 (1.411) |
| R<sub>pim</sub> | 0.037 (0.465) | 0.038 (0.497) | 0.038 (0.463) | 0.035 (0.406) | 0.054 (0.511) | 0.033 (0.532) |
| I/σ(I) | 10.3 (1.5) | 11.2 (1.7) | 11.4 (1.7) | 10.9 (1.6) | 9.1 (1.8) | 12.0 (1.7) |
| Completeness (%) |  |  |  |  |  |  |
| spherical | 62.7 (13.9) | 62.8 (13.7) | 61.4 (10.8) | 63.8 (11.1) | 53.9 (9.1) | 70.2 (16.3) |
| ellipsoidal | 83.5 (54.2) | 92.5 (59.0) | 91.1 (59.7) | 92.6 (71.0) | 89.5 (61.1) | 93.1 (61.6) |
| CC<sub>1/2</sub> | 0.997 (0.658) | 0.997 (0.581) | 0.998 (0.516) | 0.998 (0.604) | 0.997 (0.599) | 0.998 (0.573) |
| **Refinement** |  |  |  |  |  |  |
| Resolution (Å) | 61.10-2.04 | 78.94-2.12 | 54.29-2.18 | 94.63-2.15 | 93.16-2.34 | 54.93-2.05 |
| No. reflections | 16378 | 57538 | 51311 | 56713 | 35059 | 72891 |
| R<sub>work</sub> / R<sub>free</sub> (%)<sup>a</sup> | 21.8/25.5 | 22.4/24.8 | 20.9/24.2 | 22.2/24.9 | 25.5/29.5 | 22.0/25.3 |
| No. non-H atoms | 3498 | 8351 | 8375 | 8203 | 8268 | 8334 |
| Protein | 3427 | 7018 | 7072 | 6956 | 7028 | 6943 |
| DNA | 0 | 1189 | 1189 | 1189 | 1189 | 1189 |
| Metal ions | 4 | 8 | 8 | 8 | 8 | 8 |
| Water | 67 | 136 | 106 | 50 | 43 | 194 |
| B-factors |  |  |  |  |  |  |
| Protein | 38.4 | 51.7 | 60.1 | 60.4 | 64.0 | 51.6 |
| DNA | - | 60.5 | 67.1 | 71.6 | 77.9 | 74.5 |
| Metal ions | 21.1 | 38.2 | 51.8 | 44.7 | 46.7 | 39.6 |
| Water | 30.6 | 40.6 | 47.8 | 45.4 | 36.9 | 48.1 |
| All atoms | 35.8 | 49.8 | 57.5 | 57.7 | 61.7 | 51.6 |
| R.m.s. deviations |  |  |  |  |  |  |
| Bond lengths (Å) | 0.0023 | 0.0023 | 0.0021 | 0.0028 | 0.0022 | 0.0023 |
| Bond angles (°) | 1.183 | 1.120 | 1.102 | 1.158 | 1.103 | 1.125 |
| Rotamers favored/poor<br>(%)<sup>b</sup> | 96.8/0.5 | 95.7/0.5 | 95.7/0.7 | 94.6/0.5 | 95.7/0.8 | 96.5/0.4 |
| Ramachandran<br>favored/outliers (%)<sup>b</sup> | 97.2/0.0 | 96.8/0.1 | 97.0/0.2 | 96.1/0.1 | 94.8/0.1 | 98.1/0.1 |
| Rama Z-score<sup>b</sup> | -2.13 ± 0.36 | -1.57 ± 0.24 | -1.60 ± 0.23 | -1.83 ± 0.24 | -3.64 ± 0.22 | -1.55 ± 0.23 |
| Clashscore<sup>b</sup> | 1.44 | 0.52 | 0.65 | 1.38 | 1.75 | 0.72 |
| MolProbity score<sup>b</sup> | 1.03 | 0.87 | 0.88 | 1.13 | 1.29 | 0.73 |

Values in parentheses are for the highest resolution shell. Friedel pairs were merged. <sup>a</sup> R<sub>free</sub> is calculated from a randomly selected subset of ~5% of reflections excluded from refinement. <sup>b</sup> Geometry statistics were calculated with MolProbity (Williams *et al*, 2018).

We then co-crystallized IdeR with a 30-bp double-stranded DNA oligomer containing the consensus sequence TTAGGTTAGSCTAACCTAA. Crystal structures were obtained for the complexes with the physiological activator Fe^2+^ as well as the mimic Co^2+^ (Table 2). Both crystallized in space group C_2_, containing four polypeptide chains forming two dimers and two DNA strands forming a distorted B-type double helix in the asymmetric unit (Figs 2A, EV1B and EV3A). Each IdeR subunit binds to one of the four CCTAA repeats of the recognition sequence, one dimer interacting with repeats 1 and 3 and the other with repeats 2 and 4 (see Table 1). Both metal-binding sites of each IdeR subunit are occupied in both the Fe^2+^ and Co^2+^ complexes (Figs 2B and EV2B and C). The structures of these IdeR dimers are very similar to the DNA-free dimer structure (Appendix Table S1). DNA binding primarily causes a slight shift of the recognition helices, which is necessary to allow these helices to insert into the major grooves of the DNA (Fig EV1A). No significant differences between the Fe^2+^-activated and Co^2+^-activated IdeR-DNA complexes can be discerned, neither globally, nor at the metal-binding sites (Figs EV1B and EV2B and C, Appendix Table S1). Interestingly, in the DNA complex structures we observe a swap of the SH3-like domains of one subunit of each IdeR dimer with a symmetry-related chain (Fig EV3B). While we assume that it is a crystallization artifact, we cannot exclude the possibility that the SH3 domain swap is used *in vivo* to cross-link neighboring promoters.

### IdeR recognizes half binding sites

Both DNA sequences evaluated above (Fig 1) have a conserved IdeR binding site, with only three and two mismatches compared to the perfect palindromic consensus, respectively (Table 1). However, some of the predicted targets collected in Table EV1, such as C23S1, diverge more from the full consensus. The binding site at C23S1 is predicted to control the expression of the complex I NADH dehydrogenase of the respiratory chain. This sequence has 6 mismatches with the full consensus, and the distribution of those mismatches suggests that it can only be recognized by one IdeR dimer, instead of the typical two dimers (Table 1).

To date, all described IdeR/DtxR complexes with DNA involve two dimers bound to the 19-bp target sequence (Wisedchaisri *et al*, 2007; Kurthkoti *et al*, 2015; Ghosh *et al*, 2015; Wisedchaisri *et al*, 2004; Pohl *et al*, 1999). To test if the binding of only one dimer is possible with only half of the DNA target, we designed a DNA sequence harboring the two CCTAA motifs that should be recognized by one of the dimers while disrupting the sequence that should be bound by the second dimer. As can be seen in Fig 3A, IdeR is able to bind to this half binding site with an estimated *K_D_* of 78 nM, comparable to the affinity of IdeR for the previously tested C10S1 and C10S2 DNA targets (Table 1). A comparison of the electrophoretic mobility of IdeR complexes with a complete DNA target and with the half binding site clearly shows that only one IdeR dimer is bound to the half site target (Fig 3B).

**Figure 3.**
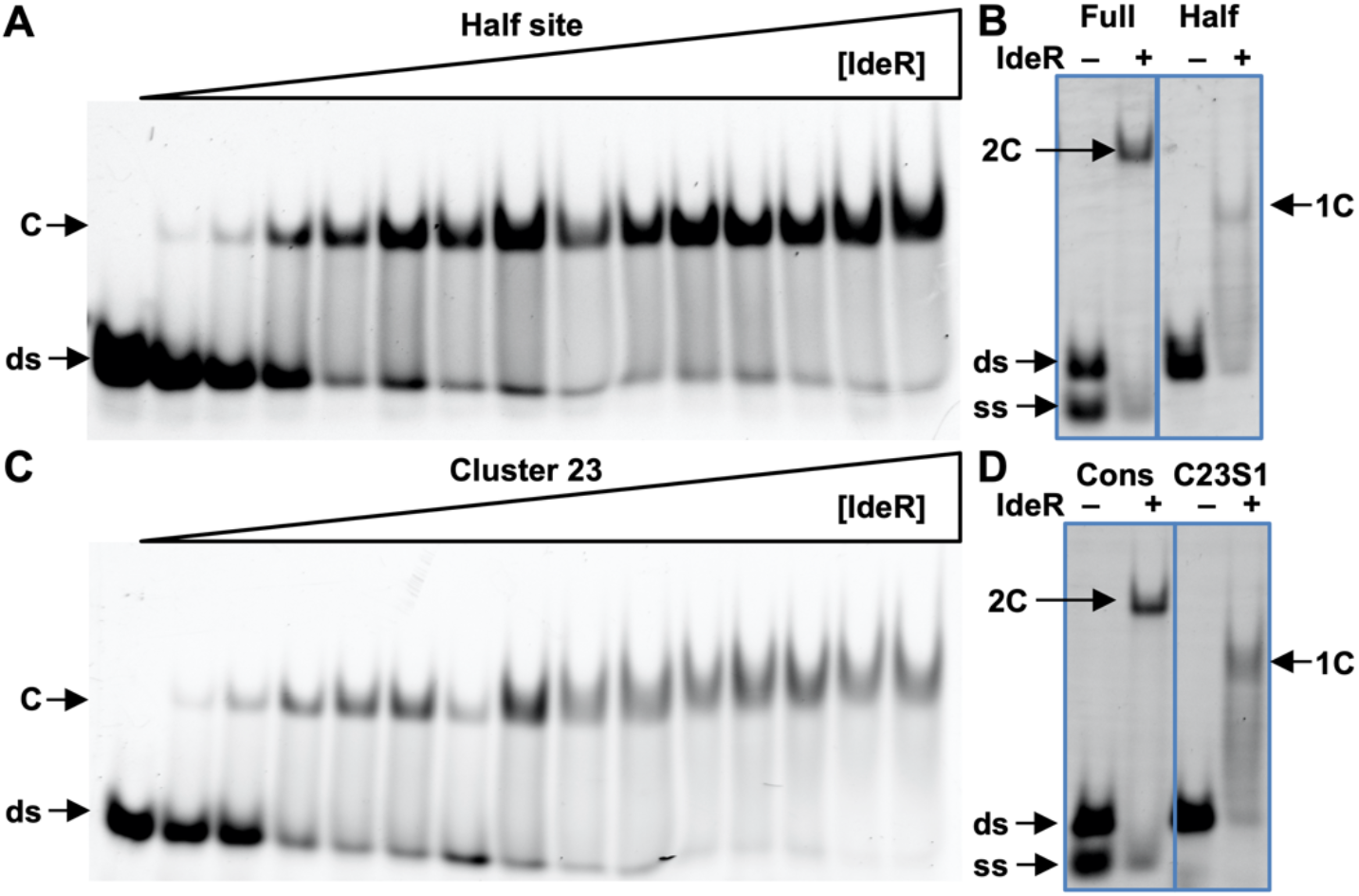
EMSA analysis of the stoichiometrical requirement of IdeR to bind its target DNA. A Binding of IdeR to a DNA sequence designed to bind only one IdeR dimer. B Binding of IdeR to the full consensus DNA sequence and the designed half binding site. C Binding of IdeR to site 1 of cluster 23. D Binding of IdeR to the full consensus DNA sequence and site 1 of cluster 23. IdeR was added to 30 nM fluorescence-labeled double-stranded DNA probe in increasing concentrations (15 nM - 2.25 μM dimer) (A, C) or at 1.5 μM (B, D) in the presence of 30 μM Co^2+^ (A, C) or 40 μM Co^2+^ (B, D) and competitor DNA. Protein-DNA complexes were resolved on a 4% Trisacetate polyacrylamide gel. The left-most lane in panels A and C is a control reaction without protein. The gel pictures in panels B and D have been edited for easier comparison between both samples. ds, unbound double-stranded DNA probe; ss, non-hybridized single-stranded DNA; C, protein-dsDNA complex; 2C, protein-dsDNA complex containing two IdeR dimers; 1C, protein-DNA complex containing one IdeR dimer.

IdeR also recognizes the C23S1 DNA target with an estimated *K_D_* of 102 nM (Fig 3C, Table 1). As expected from the sequence analysis, the binding stoichiometry of this complex is of only one dimer per DNA molecule, forming a complex similar to that observed with the half binding site target (Fig 3D). These results indicate that the DNA targets of IdeR do not require to be recognized by two dimers, and expand the number of putative targets beyond what was previously predicted for this family of regulators.

### IdeR forms very few interactions with DNA bases in the recognition sequence

*Se*IdeR interacts with DNA in the manner typical for HTH DNA-binding domains and similarly to other IdeR/DtxR-DNA complexes (Pohl *et al*, 1999; Wisedchaisri *et al*, 2004). Each IdeR monomer recognizes one of the four five-nucleotide repetitions (CCTAA) conserved in the palindromic 19-bp consensus. The HTH motif is anchored to the DNA on both sides of the major groove by hydrogen bonds with the phosphate backbone of the DNA, facilitated by residues from the first helix of the HTH motif on one side and residues from the second, so-called recognition helix on the other, and the recognition helix is thereby inserted into the major groove (Fig 4A). Notably, only one hydrogen bond interaction between the protein and a DNA base can be observed, between Gln43 and the first cytosine of each CCTAA repeat, or guanine at the central G-C basepair of the palindrome (Fig 4A and B). However, a water-mediated hydrogen bond between Gln43, the backbone carboxyl group of Pro39 and the second cytosine can also be observed in most IdeR subunits and is likely always present. Additionally, the crystal structures reveal van der Waals (vdW) interactions between Pro39 and Ser37 and the T in the third position of the repeat (Fig 4A and B). The sidechain of Pro39 is also in close proximity to the A-T basepair in the fourth position, though these vdW interactions appear to be unspecific (see below).

**Figure 4.**
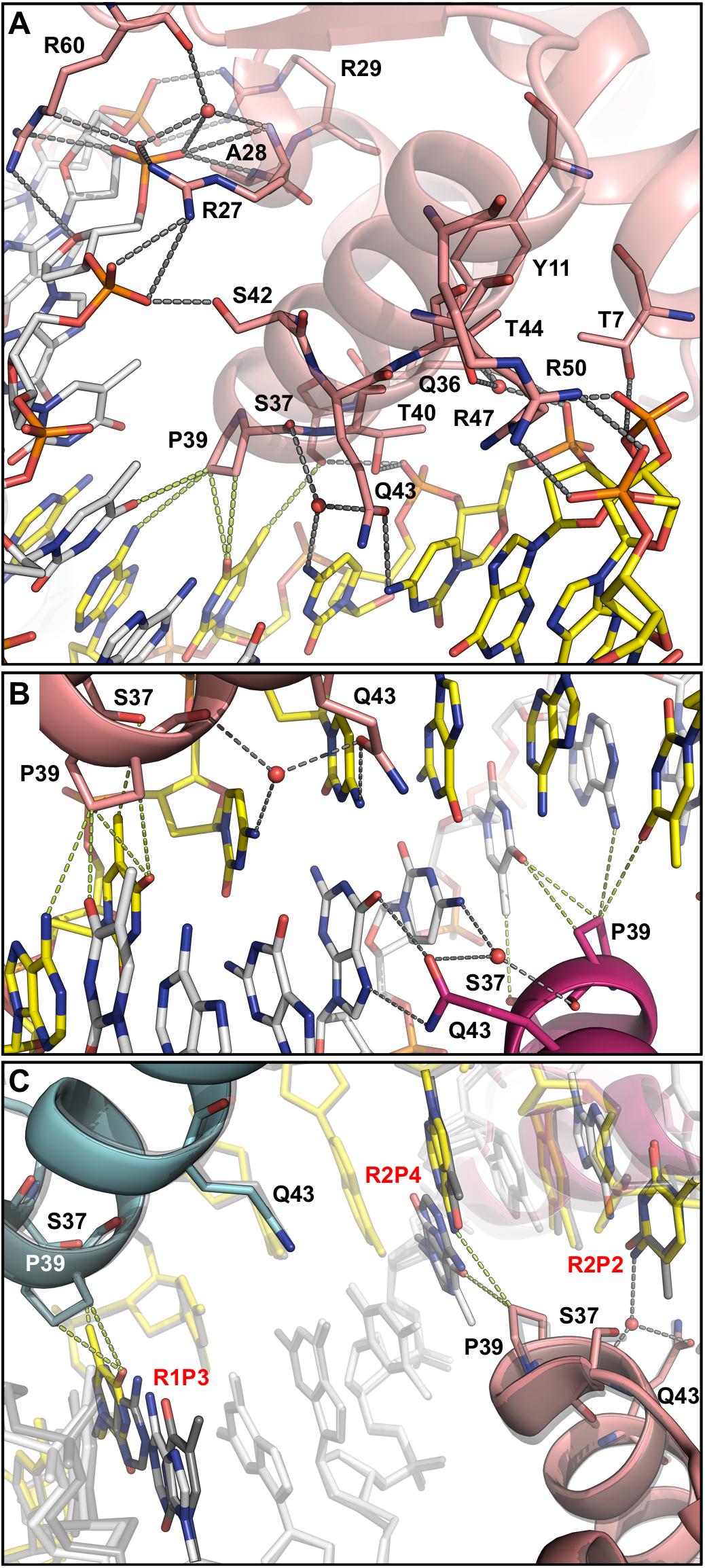
Interactions between IdeR and DNA. A Interactions between IdeR and the consensus DNA, illustrated with IdeR subunit B of the Fe^2+^-activated IdeR-consensus DNA complex. In every subunit, the side chains of Thr7, Arg27, Arg29, Gln36, Ser37, Thr40, Ser42, Arg47, Arg50, and Arg60, as well as the peptide bond amide group of Ala28 form hydrogen bonds with the phosphate backbone of the DNA. Additionally, water-mediated interactions with phosphates are formed by the side chains of Tyr11, Gln36 and Thr44, as well as the backbone amide of Arg27 and the backbone carboxyl group of Arg60. Gln43 forms a hydrogen bond with the first cytosine in the CCTAA repeat of the palindromic recognition sequence as well as a water-mediated hydrogen bond with the second cytosine. Pro39 and Ser37 form vdW interactions with the thymine in position 3 and the fourth basepair in the repeat. B Interactions between Fe^2+^-activated IdeR and DNA bases, focused on the central G-C basepair in the consensus DNA-binding sequence, which interacts with two IdeR subunits from different dimers. C Comparison of the interactions between IdeR and the DNA that are affected by the differences between the consensus sequence and the C10S1 sequence. The Fe^2+^-activated IdeR-consensus DNA complex is shown colored by subunit, while the Co^2+^-activated IdeR-C10S1 DNA complex is shown in transparent dark grey. For clarity, the DNA strands in both complexes are shown partially transparent, except for the bases that differ between the two DNA sequences, which are also colored by element. Ser37, Pro39 and Gln43 of the IdeR subunits that interact with the differing bases are shown as sticks. Interactions that are affected by the sequence differences are shown only for the consensus sequence. The C10S1 sequence differs from the consensus in the third position of the first repeat (R1P3, A instead of T), and the second and fourth position of the second repeat (R2P2, T instead of C, and R2P4, T instead of A). Hydrogen bonds are indicated by dashed grey lines, vdW interactions (distances between 3.4 - 3.7 Å) by dashed green lines.

We also determined the crystal structure of the Co^2+^-activated complex of IdeR and C10S1 DNA (Table 2). The structure of this complex does not display any significant differences compared to the complexes with the consensus sequence (Figs EV1B and EV2D, Appendix Table S1). The C10S1 sequence differs from the consensus in position 3 of the first repeat, and positions 2 and 4 of the second repeat (Table 1). These differences affect the interactions with one subunit of each IdeR dimer. Specifically, the vdW interactions with Pro39 and Ser37 of one IdeR subunit, and the vdW interactions with Pro39 of the IdeR subunit bound to the neighboring major groove are affected. However, the only notable difference regarding these distance-dependent interactions is the absence of the thymine methyl group in the first repeat, as the distances between Pro39 and the fourth base pair are very similar regardless of the nature of the bases (Fig 4C). The water-mediated hydrogen bond with Glu43 and Pro39 should not be affected by the different base in position 2 of the second repeat, but the water molecule, though likely present, was not clearly observed in the electron density and was not modelled. Despite these differences, IdeR binds to the C10S1 sequence in exactly the same way as to the consensus sequence (Fig 4C).

### DNA mutants suggest a reexamination of the role of the base interactions of IdeR

The scarcity and lack of specificity of the interactions between IdeR and DNA bases observed in the crystal structures led us to examine the mechanism of DNA recognition in more detail. To assess the role of each of the DNA base interactions with IdeR, we designed a set of DNA sequences diverging from the 19-bp consensus at different key positions. As shown in Fig 5A, IdeR seems to have a higher affinity (with a *K_D_* of ~45 nM, Table 1) for this consensus DNA sequence compared to the native binding sites tested before. The observed differences in affinity confirm the relevance of the mismatches present in the native binding sites, as some of those mismatches are located in the regions that are contacted by the recognition helix of IdeR.

**Figure 5.**
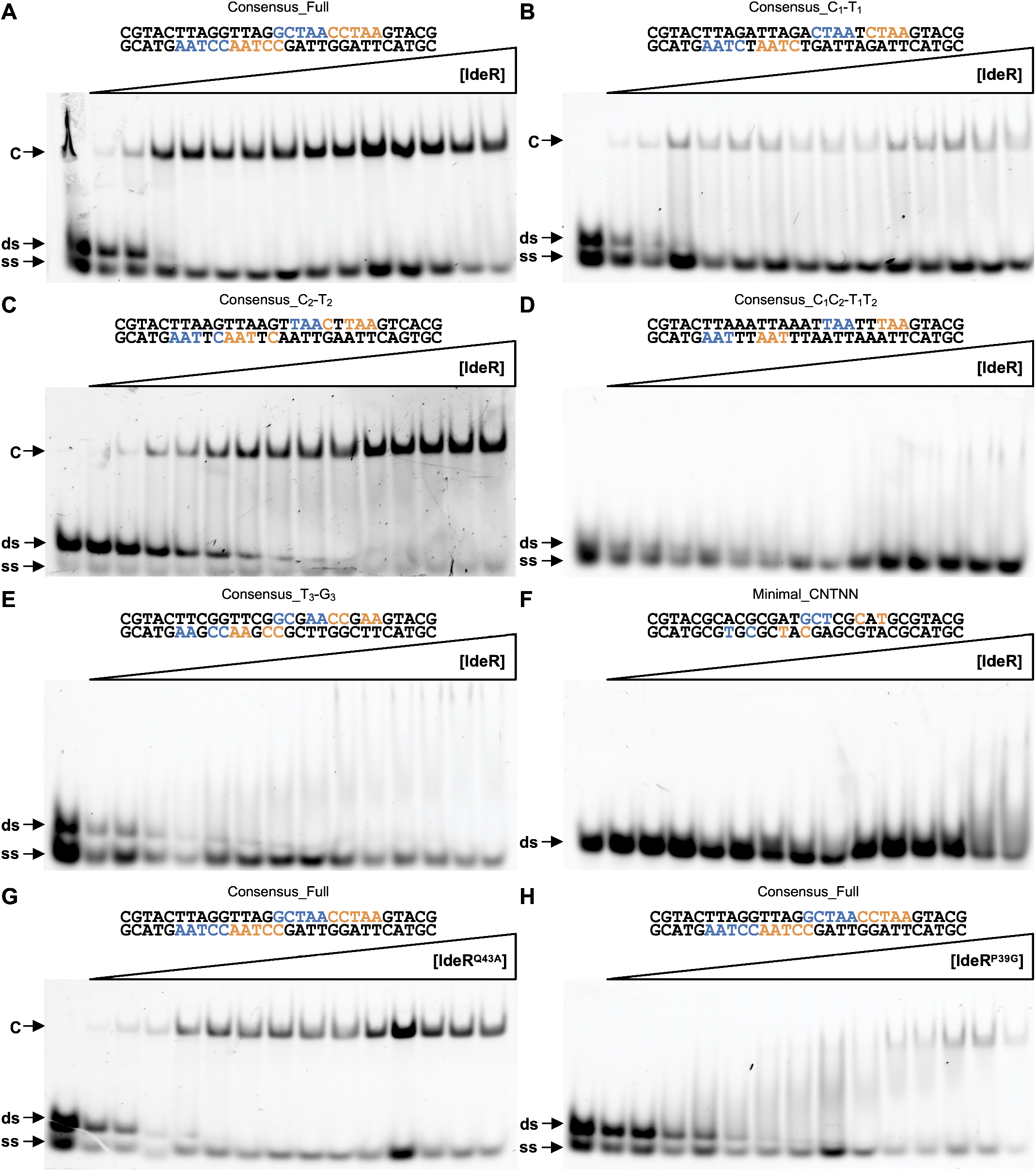
EMSA analysis of the effect of different mutations on DNA binding by IdeR. A Binding of IdeR^WT^ to the full consensus DNA sequence. B Binding of IdeR^WT^ to the Consensus_C_1_-T_1_ DNA sequence. C Binding of IdeR^WT^ to the Consensus_C_2_-T_2_ DNA sequence. D Binding of IdeR^WT^ to the Consensus_C_1_C_2_-T_1_T_2_ DNA sequence. E Binding of IdeR^WT^ to the Consensus_T_3_-G_3_ DNA sequence. F Binding of IdeR^WT^ to the Minimal_CNTNN DNA sequence. G Binding of the IdeR^Q43A^ variant to the consensus DNA sequence. H Binding of the IdeR^P39G^ variant to the consensus DNA sequence. IdeR was added in increasing concentrations (15 nM - 2.25 μM dimer) to 30 nM fluorescence-labeled doublestranded DNA probe in the presence of 30 μM Co^2+^ and competitor DNA. Protein-DNA complexes were resolved on a 4% Tris-acetate polyacrylamide gel. The left-most lane is a control reaction without protein. ds, unbound double-stranded DNA probe; ss, non-hybridized single-stranded DNA; C, protein-dsDNA complex.

The strongest base interactions of IdeR with its DNA target are represented by the hydrogen bonds between Gln43 and the first and second cytosine of each CCTAA repeat. When either of these two cytosines were mutated to a thymine on every repeat of the consensus sequence (see Consensus_C_1_-T_1_ and Consensus_C_2_-T_2_ in Table 1), no significant effect on IdeR affinity for these modified targets could be observed (Fig 5B and C). Neither of the mutations prevented IdeR recognition of the DNA targets, with no significant change in the affinity for the Consensus_C_1_-T_1_ mutant (with a *K_D_* of ~70 nM), and only a mild decrease in affinity for the Consensus_C_2_-T_2_ DNA target (with a *K_D_* of ~105 nM). However, when both cytosines were simultaneously changed to thymines (see Consensus_C_1_C_2_-T_1_T_2_ in Table 1), we did not observe any specific binding of IdeR (Fig 5D), even when increasing the IdeR concentration 10-fold compared to the conditions tested in all previous EMSAs (Fig EV4A), implying that at least one of the cytosine bases is required for IdeR recognition.

Next we replaced the thymine in position 3, which forms vdW interactions with Ser37 and Pro39 of IdeR, with a guanine in each CCTAA repeat (see Consensus_T_3_-G_3_ in Table 1). IdeR was not able to bind specifically to this DNA sequence (Fig 5E). As with the C_1_C_2_-T_1_T_2_ mutation, higher concentrations of IdeR were tested to confirm the absence of specific binding (Fig EV4B). This result suggests that a thymine in position 3 of the repeat plays a key role in IdeR recognition.

The two remaining adenines of each CCTAA repeat were also mutated to cytosine or guanine (see Consensus_A_4_A_5_-S_4_S_5_ in Table 1). Although these mutations do not affect specific base interactions with IdeR, we observed a slight drop in IdeR affinity with an estimated *K_D_* of 128 nM (Fig EV4C).

Concluding that each DNA quintet recognized by IdeR requires at minimum a thymine in the third position, and a cytosine at the first or second position, we designed a DNA sequence that should fulfill these base contact requirements for IdeR recognition, but preserves none of the other conserved bases of the recognition sequence (see Minimal_CNTNN in Table 1). However, no binding of IdeR was observed (Fig 5F) even when using high concentrations of IdeR (Fig EV4D). Noting that the number of specific base interactions provided by this sequence should not be different from those of the Consensus_C_1_-T_1_ and Consensus_C_2_-T_2_ targets tested previously (Fig 5B and C), these results indicate that a sequence-dependent recognition mechanism other than base-specific interactions plays a key role in this process.

### IdeR variants suggest an indirect readout mechanism for specific DNA binding

To clarify the relevance of the base interactions with the recognition helix of IdeR, we generated two IdeR variants, IdeR^Q43A^ and IdeR^P39G^, that should disrupt these interactions. When testing the affinity of IdeR^Q43A^ for the consensus sequence with EMSA, we did not observe any significant differences compared to IdeR^WT^ (Fig 5G; *K_D_* ~65 nM), demonstrating that the hydrogen bonds between Gln43 and the DNA bases are not important for the recognition of the DNA target. Furthermore, these results show that the absence of IdeR^WT^ binding to the sequence Consensus_C_1_C_2_-T_1_T_2_ (Fig 5D) was not due to a disruption of the Gln43-C_1_/C_2_ interactions.

To discard the possibility that other residues have taken the role of Gln43 in this IdeR variant and established new hydrogen bonds, we obtained the crystal structure of DNA-bound IdeR^Q43A^ (Table 2, Figs 6A and EV5A-C). The structure is essentially identical to that of the IdeR^WT^-DNA complexes, despite the loss of both the direct hydrogen bond between Gln43 and the first cytosine of each CCTAA repeat as well as the water-mediated hydrogen bond with the second cytosine. It is unclear if the water molecule bound to the second cytosine is lost due to the mutation or not observed as a result of the lower resolution of this structure (Table 2), but it is clearly present in one of the four IdeR^Q43A^ subunits. No additional interactions were observed in the crystal structure of this complex (Fig 6A), corroborating that the base contacts of Gln43 in the recognition helix of IdeR are not required for correct target recognition and binding.

**Figure 6.**
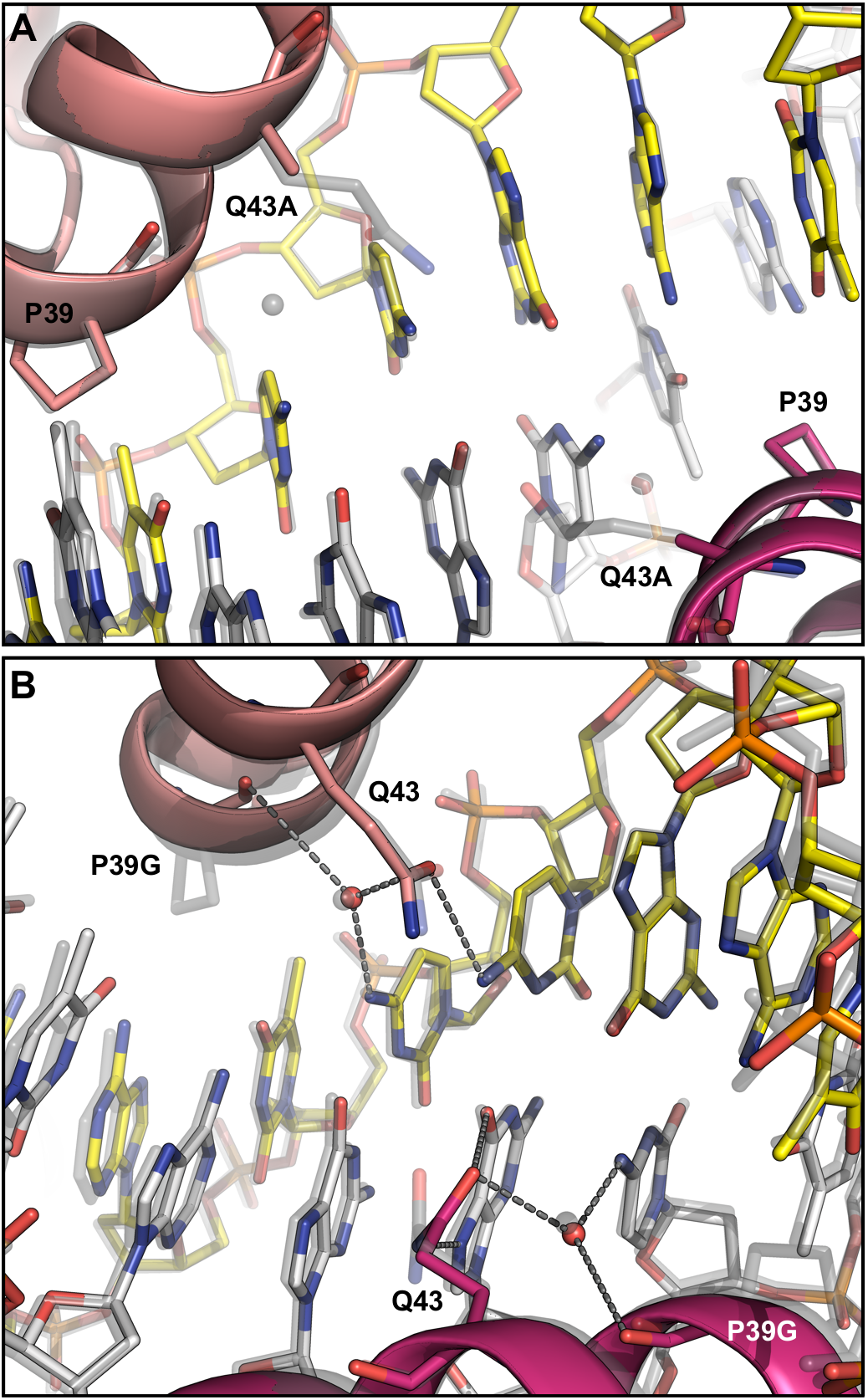
Crystal structures of IdeR variants in complex with the consensus DNA-binding sequence. A, B Interactions between Co^2+^-activated IdeR^Q43A^ (A) or IdeR^P39G^ (B) and DNA bases, focused on the central G-C basepair in the consensus DNA-binding sequence. The IdeR variants (colored by subunit) are shown superimposed with the Fe^2+^-activated IdeR^WT^-consensus DNA complex structure (transparent grey). In B, hydrogen bonds in the IdeR^P39G^ structure are indicated by dashed grey lines.

As for the IdeR^P39G^ variant, the EMSA results show specific binding to the consensus target, but the affinity is significantly affected by the mutation, resulting in a *K_D_* of ~264 nM (Fig 5H). Due to the special structural features of both proline and glycine residues we cannot conclude whether this loss of affinity is caused by the absence of the vdW interactions with the DNA, or if this protein variant is functionally impaired. However, as IdeR^P39G^ is still able to recognize its target and form a stable complex, we can reason that the absence of IdeR^WT^ binding to the Consensus_T_3_-G_3_ sequence (Fig 5E) was not caused by the disruption of the vdW interactions between Pro39 and the thymine in position 3. We also obtained the crystal structure of IdeR^P39G^ bound to the consensus DNA to confirm that no other interactions with the essential thymines of the recognition sequence are formed in this IdeR variant (Table 2, Figs 6B and EV5). Despite the drastic difference between the wild-type and mutated residue, in the DNA-bound state the structure of the recognition helix is essentially unaffected by the mutation. The mutation does not disrupt the hydrogen bond formed by the backbone carboxyl group of residue 39 with the water molecule which is also bound to Gln43 and the cytosine in position 2 of the repeat.

Altogether, these results provide evidence of an indirect readout mechanism for IdeR, where the protein perceives the sequence-dependent structure of the DNA by interacting with its phosphate backbone, instead of directly contacting the specific bases in the DNA. This mechanism also agrees with the cooperativity observed for the interaction. The absence of one-dimer complexes in any of the EMSAs performed with all full-target DNA sequences indicates that the binding of both IdeR dimers to the full DNA sequence is cooperative. As we observe no interaction between these two protein dimers in the crystal structures, the cooperativity must arise from a structural change of the DNA double helix upon IdeR binding.

## DISCUSSION

*Se*IdeR recognizes the 19-bp DNA consensus sequence established for other IdeR/DtxR iron sensors consisting of a palindromic repeat of four CCTAA motifs. Each of these four repeats is recognized by an IdeR monomer with its HTH DNA-binding motif, resulting in a two-dimer complex with the full DNA sequence. This sequence can be found at 37 regions in the *S. erythraea* genome with different degrees of conservation. The number of targets is consistent with those found in other organisms such as *M. tuberculosis* or *C. diphtheriae*. IdeR binding was confirmed for the siderophore cluster comprised of genes SACE_2689 to SACE_2697 and for the cluster coding for the respiratory chain NADH dehydrogenase Nuo complex I (genes SACE_6902 to SACE_6889).

Complex I of the respiratory chain has been linked to ROS production in mitochondria (Imlay, 2013). Little is known about complex I contribution to ROS formation in bacteria, but some species seem to favor the use of complex II in high oxygen conditions, despite the fact that complex II has a bigger role in ROS production in these species (Wackwitz *et al*, 1999; Messner & Imlay, 1999). The nuo gene cluster has been shown to be induced by iron in other bacteria such as *M. tuberculosis* and *Geobacter sulfurreducens* (Rodriguez *et al*, 2002; Embree *et al*, 2014). While no iron-dependent regulator has been identified as responsible for this induction in *M. tuberculosis*, in *G. sulfurreducens* it was found to be under the control of the ferric uptake regulator Fur (Rodriguez *et al*, 2002; Embree *et al*, 2014). Our results indicate that in *S. erythraea*, complex I production is controlled by IdeR.

The binding of IdeR to the promoter of the nuo gene cluster in *S. erythraea* is the first reported example of the formation of an IdeR-DNA complex with only one dimer of the transcriptional regulator. This finding suggests a redefinition of the consensus of the DNA targets for iron-dependent IdeR/DtxR regulators from the 19-bp TTAGGTTAGSCTAACCTAA to the 10-bp TTAGGNNNNCCTAA consensus, which may result in the discovery of new targets in other IdeR/DtxR species. What the function of IdeR at such half sites is remains to be investigated.

In this work, we show evidence of an indirect readout mechanism for IdeR. Indirect readout is a recognition mechanism based on the structural reading, rather than molecular reading, of a DNA sequence by its DNA-binding protein (Abe *et al*, 2015; Yang *et al*, 2017; Dorman & Dorman, 2017). Although the concept of indirect readout is well established among eukaryotic transcription factors, it is frequently overlooked and underestimated for prokaryotic transcription factors (Dorman & Dorman, 2017; Sarvan *et al*, 2018). This recognition mechanism was previously proposed for DtxR-like proteins by Lee and Holmes (Lee & Holmes, 2000), who considered that the Gln43 interactions with the cytosine bases were not enough to explain the specificity of this transcription factor. However, Chen *et al.* (Chen *et al*, 2000) described the vdW interactions with the thymine bases, which would theoretically add to the Gln43 interactions to support a direct readout of the DNA sequence for this type of proteins. Here we show that neither of these interactions determines the recognition process, and that an indirect readout mechanism is required to explain DNA recognition by IdeR/DtxR-like proteins.

It might be argued that the vdW interactions with the thymine bases, mediated by Pro39 and Ser37, are still relevant for DNA recognition, as the P39G variant did not show the same affinity for the consensus sequence as IdeR^WT^. However, several pieces of evidence suggest that these interactions are nonessential: (1) considering that the Gln43 interactions are not required for recognition, the strength of the vdW interactions does not suffice to account for specific recognition of the full target sequence; (2) the IdeR^P39G^ variant is still able to specifically recognize its DNA target, implying a recognition mechanism independent of these vdW interactions; (3) the loss of affinity of this IdeR variant may be caused by the biochemical nature of the exchanged residues that might result in undesired effects on the dynamics of the protein, affecting the flexibility of the HTH motif, and thereby the affinity for its DNA target; (4) as shown by the lack of binding of IdeR to the Minimal_CNTNN DNA sequence, the presence of all thymine bases involved in the vdW interactions is not enough for target recognition; (5) the work done by Chen *et al.* (Chen *et al*, 2000) and Spiering *et al.* (Spiering *et al*, 2003), although highlighting the relevance of the thymine bases for recognition, showed binding of two dimers to a DNA sequence lacking both thymine bases for one of those dimers, again implying a recognition mechanism that is independent of the vdW interactions with the thymines.

In conclusion, our results indicate that IdeR recognizes its targets by reading the sequence-dependent DNA backbone structure. The similarities of the HTH motifs of most iron-dependent IdeR/DtxR regulators, in line with the fact that most IdeR/DtxR proteins recognize the same consensus sequence, suggest that they use the same DNA recognition mechanism. Several sequence-based bioinformatic analyses have been performed to identify the targets of DtxR-like regulators in bacterial genomes, resulting in several dozens of different potential binding sites for this type of protein (Gold *et al*, 2001; Kunkle & Schmitt, 2003; Yellaboina *et al*, 2004; Brune *et al*, 2006; Yellaboina *et al*, 2006; Granger *et al*, 2013; Deng & Zhang, 2015; Cheng *et al*, 2018). Although our results imply that a structural approach might result in more accurate predictions for DtxR-like targets, the current bioinformatic tools do not allow such analyses on a genomic scale (Chiu *et al*, 2015). However, until such a tool is developed, the sequence-based predictions have proven to be an efficient strategy to find potential targets for iron-dependent DtxR regulators.

## MATERIAL AND METHODS

### Identification of IdeR binding sites

To identify the putative targets of IdeR in the genome of *S. erythraea*, a pattern search was performed using the Pattern Locator software developed by CMBL (https://www.cmbl.uga.edu/software/patloc.html) (Mŕazek & Xie, 2006), searching for the full 19-bp consensus sequence with a 6 mismatch allowance, as well as the half binding site with only one mismatch. The resulting hits were manually curated by discarding all non-intergenic sequences and sequences located further than 500 bp from the closest annotated starting codon. A total of 37 sequences were selected from the resulting list as likely IdeR targets, either by sequence conservation, redundancy in their gene cluster, or by predicted product.

### Cloning

The full-length *S. erythraea* IdeR (accession number WP_009947362.1) coding sequence was PCR-amplified from genomic DNA (DSM number 40517) and inserted into a modified version of pET-28a(+) (Novagen), which encodes the recognition sequence for Tobacco Etch Virus (TEV) protease instead of thrombin, using the *NdeI* and *HindIII* restriction sites (see Appendix Table S2 for primer sequences). The resulting construct was verified by DNA sequencing. It encodes full-length IdeR with a TEV-cleavable N-terminal hexahistidine tag and no C-terminal tag, so that after TEV cleavage, full-length IdeR including the N-terminal Met residue remains with two additional N-terminal amino acids (Gly-His). Point mutations (Q43A, P39G) were introduced into this construct by PCR-based site-directed mutagenesis using the QuikChange™ method and verified by DNA sequencing (see Appendix Table S2 for primer sequences).

### Protein production and purification

*E. coli* BL21(DE3) (Novagen) cells transformed with the plasmid encoding IdeR^WT^ or one of its engineered variants IdeR^Q43A^ or IdeR^P39G^ were grown in terrific broth (TB) medium supplemented with 50 μg/ml kanamycin at 37°C to an OD_600_ of ~0.5. Expression was then induced by adding 0.1 mM isopropyl-β-D-thiogalactopyranosid (IPTG) and the cultures were harvested after over-night incubation at 20°C. The harvested cells were resuspended in IdeR lysis buffer (25 mM 2-(N-morpholino)ethanesulfonic acid [MES] pH 6.0, 450 mM NaCl, 10% (v/v) glycerol) with the addition of 5 mM MgSO_4_, 1 mM phenylmethylsulfonyl fluoride (PMSF), DNase and lysozyme, and lysed with a cell disruptor (Constant Systems). The lysate was cleared by centrifugation and the supernatant was applied to a Ni^2+^-nitrilotriacetic acid (NTA) resin (Ni Sepharose 6 Fast Flow, Cytiva) gravity flow column. The column was washed with at least 10 column volumes (CV) IdeR wash buffer (25 mM MES pH 6.0, 450 mM NaCl, 60 mM imidazole, 10% (v/v) glycerol) and the protein eluted with 5 CV IdeR elution buffer (25 mM MES pH 6.0, 450 mM NaCl, 500 mM imidazole, 10% (v/v) glycerol). The eluted protein sample was then concentrated to an appropriate volume and exchanged into IdeR lysis buffer using PD10 columns (Cytiva). The His-tag of the recombinant IdeR was cleaved over-night at room temperature by adding 0.5 mM EDTA, 10 mM β-mercaptoethanol and TEV protease at a ratio of 1 μM TEV per 100 μM IdeR monomer. The digested protein sample was diluted with IdeR lysis buffer to decrease the β-mercaptoethanol concentration below 5 mM, and imidazole was added to a final concentration of 60 mM. The sample was then again applied to a Ni^2+^-NTA gravity flow column and the flow-through, containing the tag-free IdeR, collected. The column was washed with 5 CV of IdeR wash buffer and the flowthrough and wash fractions were combined and concentrated to a protein concentration of ~0.8 mM (20 mg/ml). Protein concentration was determined using a calculated extinction coefficient at 280 nm of 15.47 mM^-1^ cm^-1^ for the IdeR monomer (Gasteiger *et al*, 2005). The protein was then aliquoted, flash-frozen in liquid nitrogen and stored at −80°C until further use.

His-tagged TEV_SH_ protease (Van Den Berg *et al*, 2006) was produced and purified similarly as IdeR, with the following differences. *E. coli* BL21(DE3) (Novagen) cells transformed with the plasmid encoding TEV_SH_ (Van Den Berg *et al*, 2006) were grown in TB medium supplemented with 50 μg/ml ampicillin, and expression was induced with 1 mM IPTG. TEV lysis buffer contained 50 mM Tris-HCl pH 8.0, 200 mM NaCl and 10% (v/v) glycerol. The cleared lysate was incubated with Ni^2+^-NTA resin for 1 h at 4°C before the slurry was transferred into gravity flow columns and washed with at least 10 CV TEV wash buffer (50 mM Tris-HCl pH 8.0, 200 mM NaCl, 10% (v/v) glycerol, 60 mM imidazole). TEV protease was eluted with 5 CV TEV elution buffer (50 mM Tris-HCl pH 8.0, 200 mM NaCl, 10% (v/v) glycerol, 300 mM imidazole), concentrated to an appropriate volume, and exchanged into TEV lysis buffer using PD10 columns (Cytiva). The purified protease was then again concentrated to ~200 μM, following which EDTA was added to a final concentration of 2 mM, dithiothreitol to 5 mM, and glycerol to a final concentration of 50% (v/v), so that the final concentration of TEV protease was ~100 μM. TEV protease concentration was determined using an extinction coefficient at 280 nm of 36.13 mM^-1^ cm^-1^ (Van Den Berg *et al*, 2006). TEV protease was aliquoted, flash-frozen in liquid nitrogen and stored at −80°C until further use.

### Total-reflection X-ray fluorescence (TXRF) analysis of protein metal contents

Metal contents of IdeR protein preparations were quantified using total-reflection X-ray fluorescence (TXRF) analysis on a Bruker S2 PicoFox instrument (Klockenkämper, 1997). A gallium standard (Sigma) was added to the samples (v/v, 1:1) prior to the measurements. Technical duplicates were prepared of each sample. TXRF spectra were analyzed using the routines provided with the spectrometer. The IdeR batches as purified contained ~13% Ni and ~17% Fe.

### Preparation of double-stranded DNA

Forward and reverse DNA oligonucleotides containing the different target sequences designed for DNA-binding analysis and crystallization studies were obtained from Eurofins or Thermo Fisher Scientific with or without a 5’ Cy5 or FAM fluorescent label (see Appendix Table S2). To prepare double-stranded DNA, each forward and reverse oligonucleotide sample pair was resuspended in DNA buffer (40 mM Tris-HCl pH 7.4, 100 mM NaCl, 10 mM MgSO_4_) and mixed to a final concentration of 375 μM (for cocrystallization with IdeR) or 10 μM (for EMSA). All mixtures were then heated at 95°C for 10 min and slowly cooled down to room temperature in a thermal cycler.

### Electrophoretic mobility shift assays (EMSAs)

EMSAs were performed after mixing the fluorescently-labelled double-stranded DNA samples at 30 nM with IdeR protein samples at different concentrations. All mixtures were prepared in TAKA buffer (15 mM Tris-acetate pH 7.3, 4 mM potassium-acetate), using poly-dIdC (Poly(deoxyinosinic-deoxycytidylic acid; Sigma-Aldrich) as competitor DNA at a final concentration of 0.5 mg/ml, and incubated at room temperature for 20 min. Each EMSA was run at 4°C in a 4% native polyacrylamide gel for 30 - 40 min at 20 mA. Fluorescence of the unbound DNA and the DNA bound to IdeR was then observed in a BioRad ChemiDoc MP Imager at the proper wavelength for each of the labels. EMSAs were performed at least twice and using different protein concentrations to ensure reproducibility. For the Consensus_Full sequence, both 5’ Cy5 and 5’ FAM labels were tested to exclude effects of the label on IdeR binding (see Appendix Table S2). For the estimation of the dissociation constants (*_D_*), the band intensities in the EMSA gel images were estimated using ImageJ (https://imagej.nih.gov/ii/) (Schneider *et al*, 2012), the relative amount of bound and unbound DNA was calculated from the band intensities, and the resulting data was fit to the Hill-Langmuir equation using MATLAB.

### Crystallization and data collection

Crystallization conditions were screened using commercial kits (Molecular Dimensions) in sitting-drop vapor diffusion setups at 20°C using a Mosquito^®^ Crystal liquid handling robot (SPT Labtech), followed by optimization of the identified conditions.

To obtain the structure of IdeR^WT^ complexed with cobalt, a 12 mg/ml sample of IdeR^WT^ was mixed with 1 mM CoCl2 and 0.3 mg/ml trypsin and incubated at room temperature for 30 min. The digested sample was flash-frozen in liquid nitrogen and stored at −80°C until the next day. The protein was then crystallized in a sitting-drop vapor diffusion experiment at 20°C by mixing 110 nl of the digested sample with 90 nl 25% (w/v) PEG 1500 using the Mosquito^®^ Crystal robot. Crystals were flash-cooled in liquid nitrogen without the addition of cryoprotectant. A dataset was collected at 100 K at beamline I04 of the Diamond Light Source (Didcot, UK) (Table 2).

To obtain structures of protein-DNA complexes, a 10 mg/ml protein solution, containing 1 mM CoCl2 or Fe(NH4)2(SO4)2 and 150 μM double-stranded DNA (see Appendix Table S2), was incubated at room temperature for 10 min. For Co^2+^-activated and Fe^2+^-activated IdeR^WT^ complexed with consensus DNA, Co^2+^-activated IdeR^WT^ with C10S1 DNA, as well as the Co^2+^-activated P39G variant in complex with consensus DNA, this solution was then mixed with crystallization solution containing 30% (w/v) PEG 2000 monomethyl ether, 200 mM ammonium sulfate and 100 mM sodium acetate at pH 4.6 in a sitting-drop vapor diffusion experiment at 20°C using the Mosquito^®^ Crystal robot. The total drop volume was 200 nl and the protein volume was 67 nl, 100 nl or 133 nl. Crystals of the Co^2+^-activated IdeR^Q43A^-consensus DNA complex were obtained in a hanging-drop vapor diffusion experiment at 20°C by manually mixing 1.2 μl protein solution with 0.8 μl crystallization solution and 0.4 μl seed stock consisting of microcrystals of the same protein-DNA complex. The crystallization solution contained 29% (w/v) PEG 3350, 280 mM ammonium sulfate and 100 mM MES at pH 6.5. Crystals were flash-cooled in liquid nitrogen without the addition of cryoprotectant. All datasets of IdeR-DNA complexes were collected at 100 K at the BioMAX beamline of the MaxIV laboratory (Lund, Sweden) (Table 2).

### Structure determination, model building and refinement

All datasets were processed with the autoPROC toolbox (Vonrhein *et al*, 2011) including the STARANISO server (http://staraniso.globalphasing.org/cgi-bin/staraniso.cgi), as well as XDS (Kabsch, 2010), POINTLESS (Evans, 2006), AIMLESS (Evans & Murshudov, 2013) and other CCP4 programs (Winn *et al*, 2011). Since diffraction was significantly anisotropic in all cases, elliptical diffraction cutoffs were chosen using STARANISO based on the criterion that the local I/σ(I) ≥ 1.20. Co^2+^-activated IdeR crystallized in space group P1 with one IdeR dimer in the asymmetric unit. The structure was solved by molecular replacement using PHASER (McCoy *et al*, 2007) and chain A of the structure of *M. tuberculosis* IdeR (PDB ID 1U8R) (Wisedchaisri *et al*, 2004) as search model. All DNA complexes were in space group C_2_ with two IdeR dimers and one DNA duplex in the asymmetric unit, with an NCS rotation axis along the DNA double helix axis leading to I222 pseudosymmetry. The DNA complex structures were solved by molecular replacement using the *Se*IdeR monomer structure as a search model, and the DNA chains were manually built in Coot (Emsley *et al*, 2010). Refinement was carried out with REFMAC5 (Murshudov *et al*, 2011) and iterated with rebuilding in Coot. Refinement included bulk solvent corrections, individual atomic coordinate and isotropic *B* factor refinement. Metal-ligand bond lengths were not restrained and riding hydrogens were used during refinement. Solvent molecules were added with the “find waters” function in Coot and manually curated. Structures were validated using MolProbity (http://molprobity.biochem.duke.edu/) (Williams *et al*, 2018). Data and refinement statistics are given in Table 2. DNA conformation was analyzed using wDSSR (http://wdssr.x3dna.org/) (Lu *et al*, 2015). All figures were prepared with PyMOL (version 2.4.1; Schrödinger, LLC).

## DATA AVAILABILITY

Atomic coordinates and structure factors for the reported crystal structures have been deposited with the Protein Data Bank under accession numbers:

- 7B1V: Co^2+^-activated IdeR^WT^ (https://doi.org/10.2210/pdb7B1V/pdb)
- 7B1Y: Co^2+^-activated IdeR^WT^ in complex with consensus DNA (https://doi.org/10.2210/pdb7B1Y/pdb)
- 7B20: Fe^2+^-activated IdeR^WT^ in complex with consensus DNA (https://doi.org/10.2210/pdb7B20/pdb)
- 7B23: Co^2+^-activated IdeR^WT^ in complex with C10S1 DNA (https://doi.org/10.2210/pdb7B23/pdb)
- 7B24: Co^2+^-activated IdeR^P39G^ in complex with consensus DNA (https://doi.org/10.2210/pdb7B24/pdb)
- 7B25: Co^2+^-activated IdeR^Q43A^ in complex with consensus DNA (https://doi.org/10.2210/pdb7B25/pdb)

## ACKNOWLEDGEMENTS

We thank the staff at beamlines I04/Diamond Light Source and BioMAX/MAX IV for assistance with X-ray data collection, Martin Högbom for access to the TXRF spectrometer and Hugo Lebrette for assistance with TXRF measurements, as well as Martin Högbom and Simon Larsson for critical reading of the manuscript. DM would like to thank the tutors at the crystallographic school SEA COAST 2020 (https://seacoast.kmutt.ac.th/) for help with data processing. This work was supported by the Swedish Research Council (grant number 2016-03770), Carl Tryggers Stiftelse för Vetenskaplig Forskning (grant numbers CTS 17: 170 and CTS 19: 121) and the Faculty of Science and Technology, Uppsala University (to JJG).

## AUTHOR CONTRIBUTIONS

FJM-T performed genomic and bioinformatic analyses and DNA-binding assays. DM performed protein production and purification, mutagenesis, crystallization and structure determination. JJG performed cloning. JJG and FJM-T designed the study. All authors analyzed the data and wrote the manuscript.

## EXPANDED VIEW FIGURES

**Figure EV1.**
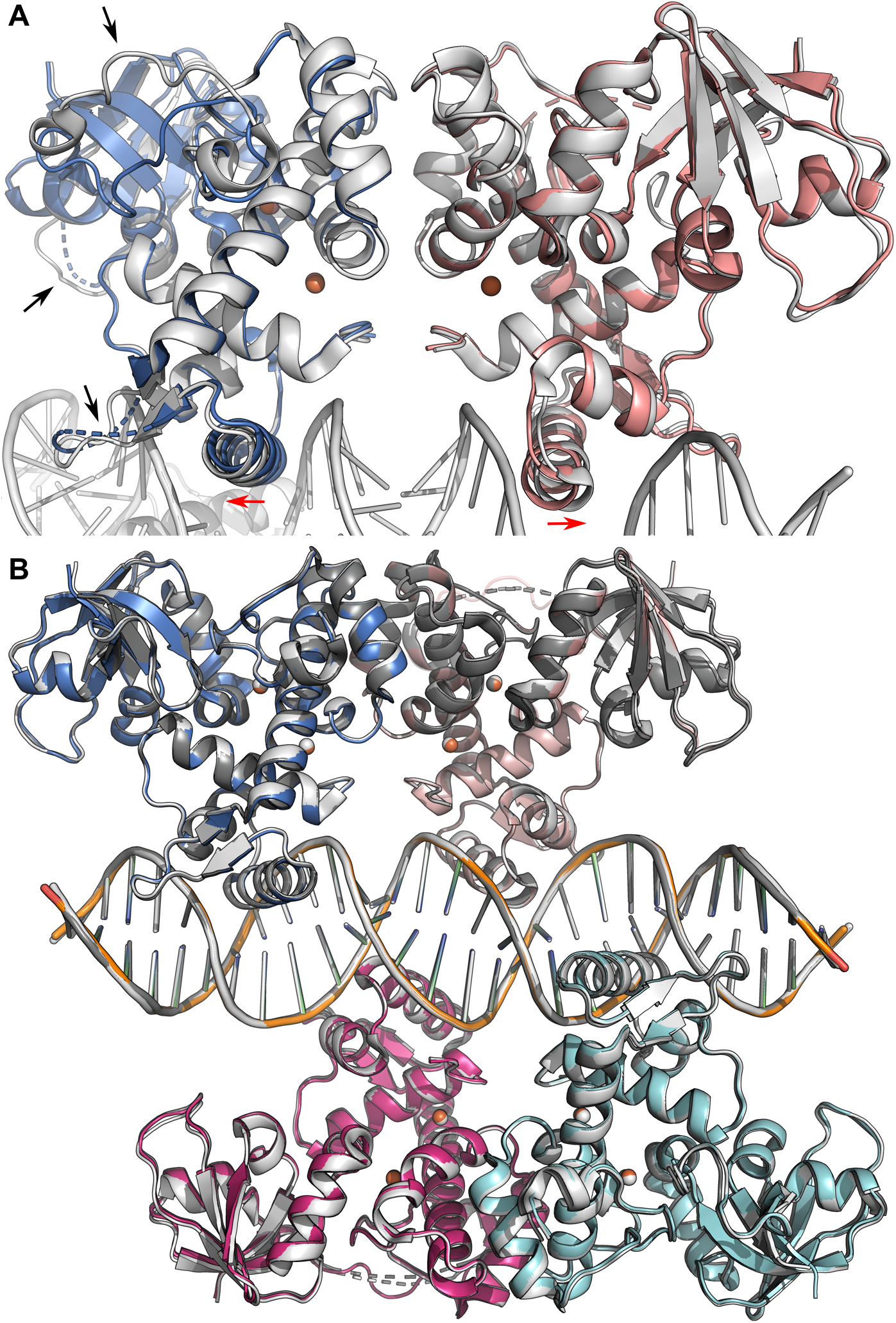
Comparison of different IdeR^WT^ complex structures. A Overall structure of the Co^2+^-activated IdeR dimer without DNA, compared to Fe^2+^-activated IdeR in complex with the consensus DNA-binding sequence. The DNA-free IdeR dimer is shown colored by subunit, superimposed with the Fe^2+^-activated IdeR-consensus DNA complex (light grey). Metal ions are shown as spheres. DNA binding causes a slight reorientation of the recognition helices (indicated by red arrows), which is required to allow the helices to insert into the major grooves of the DNA. In the DNA-bound state, several loops in the DNA-binding and SH3-like domains also assume a different or more ordered conformation (indicated by black arrows). B Superposition of the Fe^2+^-activated (colored by subunit) and Co^2+^-activated (light grey) IdeR complexes with the consensus DNA-binding sequence and the Co^2+^-activated IdeR complex with the C10S1 sequence (dark grey).

**Figure EV2.**
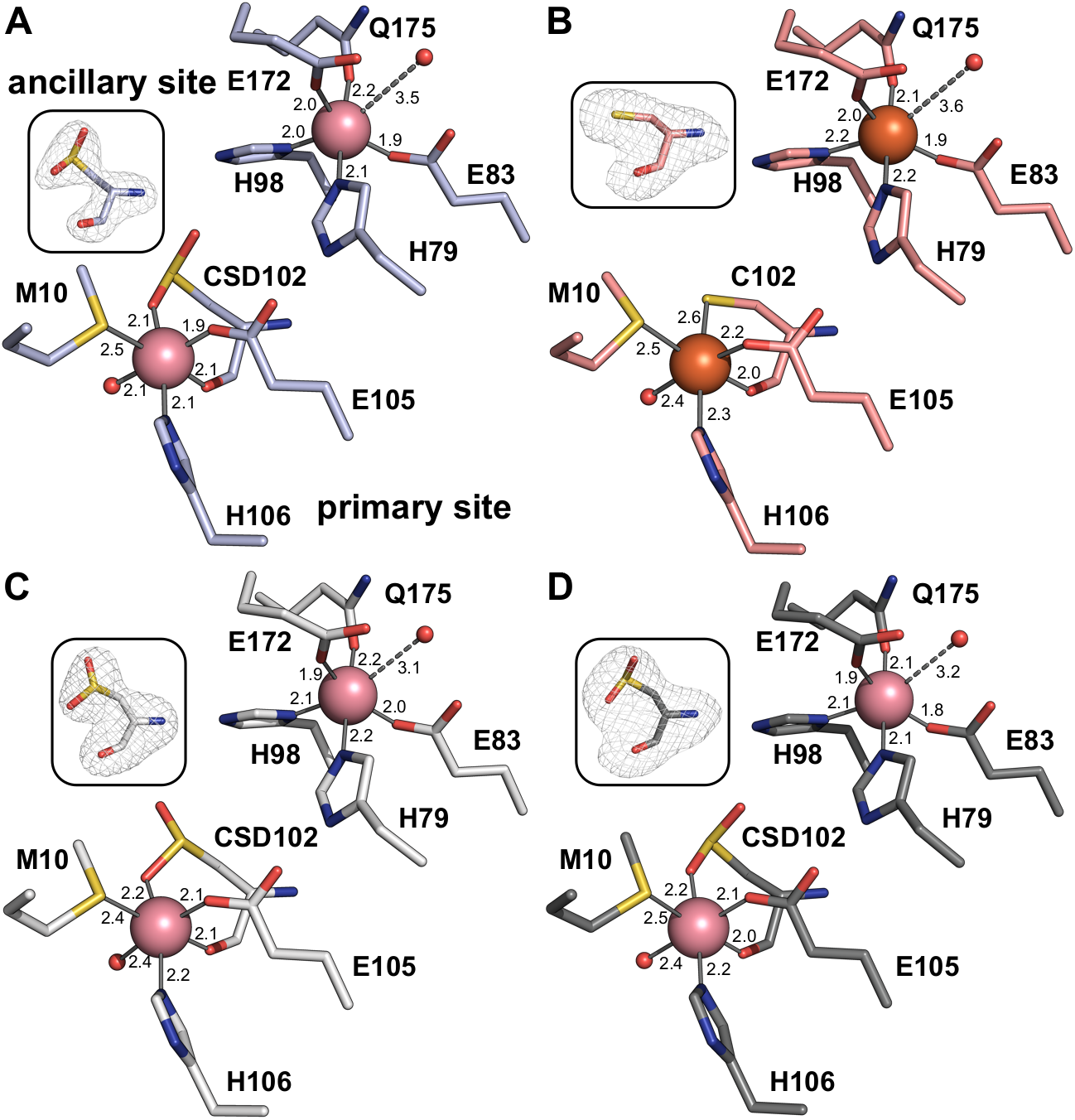
The metal-binding sites in the different IdeR^WT^ complex structures. **A** The metal-binding sites in subunit B of the Co^2+^-activated IdeR dimer without DNA. **B** The metal-binding sites in IdeR subunit B in the Fe^2+^-activated IdeR-consensus DNA complex. **C** The metal-binding sites in IdeR subunit D in the Co^2+^-activated IdeR-consensus DNA complex. **D** The metal-binding sites in IdeR subunit B in the Co^2+^-activated IdeR-C10S1 DNA complex. At the resolution of these structures, no significant differences between the complexes with the physiological activator Fe^2+^ and the mimic Co^2+^ can be discerned. Oxidation of the primary site ligand Cys102, caused by radiation damage, was observed to varying degrees in the structures described in this study. If oxidation had occurred to a significant degree, the ligand was modelled as *S*-sulfinocysteine (CSD). Metal-ligand bonds are indicated by grey lines, the dashed line between the ancillary site metal ion and water ligand indicating a long, weak bond. Bond distances are given in Å. Insets show *mF_o_-DF_c_* omit electron density for C102/CSD102 as grey mesh contoured at +4 σ.

**Figure EV3.**
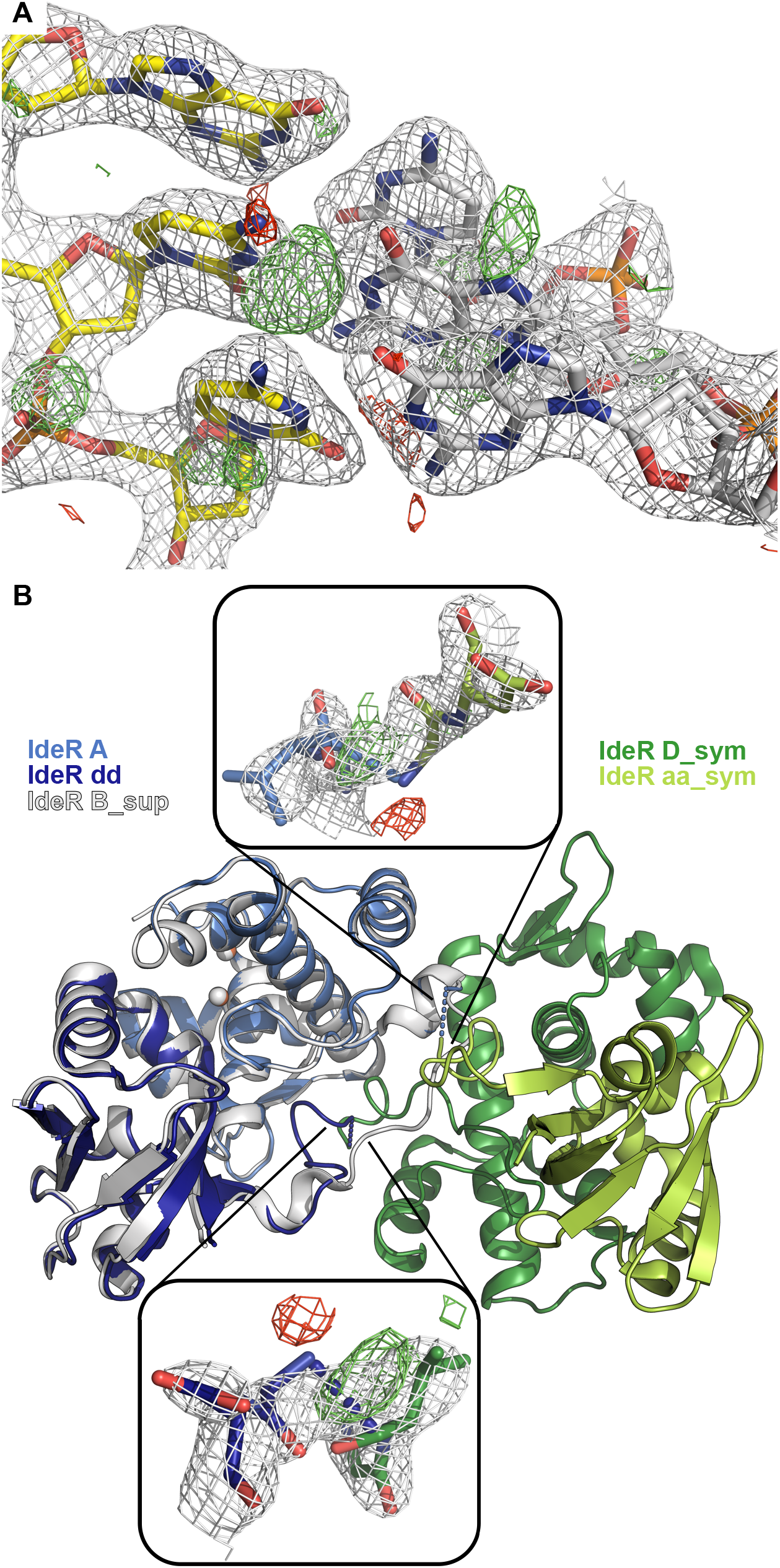
Orientation of the DNA and domain swap in crystals of IdeR-DNA complexes. A Due to the palindromic nature of the consensus DNA sequence, crystals of IdeR-consensus DNA complexes contain the DNA double helix in both orientations, as evidenced by the ambiguous *2mF_o_-DF_c_* electron density as well as the difference density at the central G-C basepair, in contrast to the well-defined electron density for the surrounding basepairs in the Fe^2+^-activated IdeR-consensus DNA complex. *2mF_o_-DF_c_* electron density is shown as grey mesh, contoured at 2 σ, *mF_o_-DF_c_* density as green mesh at +3 σ and red mesh at −3 σ. B The SH3-like domains of one subunit of each IdeR dimer in the DNA complexes are swapped with a symmetry-related chain. Due to crystal packing interactions, this domain swap is possible for IdeR chains A and D, but not chains B and C. This is illustrated here on the Fe^2+^-activated IdeR-consensus DNA complex. Shown are IdeR chain A (marine), consisting of the DNA-binding and dimerization domains, and the associated swapped SH3-like domain (chain dd, dark blue), superimposed with IdeR chain B (light grey), which represents a complete three-domain IdeR monomer in which the loop connecting the dimerization and SH3-like domains adopts a different conformation, as well as IdeR chains D_sym and aa_sym (dark green and lime green, respectively) of the symmetry mate which swaps the SH3-like domain with chain A. The connections of the A and aa_sym or D and dd_sym chains are indicated by dashed lines (marine and dark blue, respectively). Insets show *2mF_o_-DF_c_* electron density for the two residues on either side of the asymmetric unit border as grey mesh, contoured at 1.5 σ, *mF_o_-DF_c_* density as green mesh at +3 σ and red mesh at −3 σ.

**Figure EV4.**
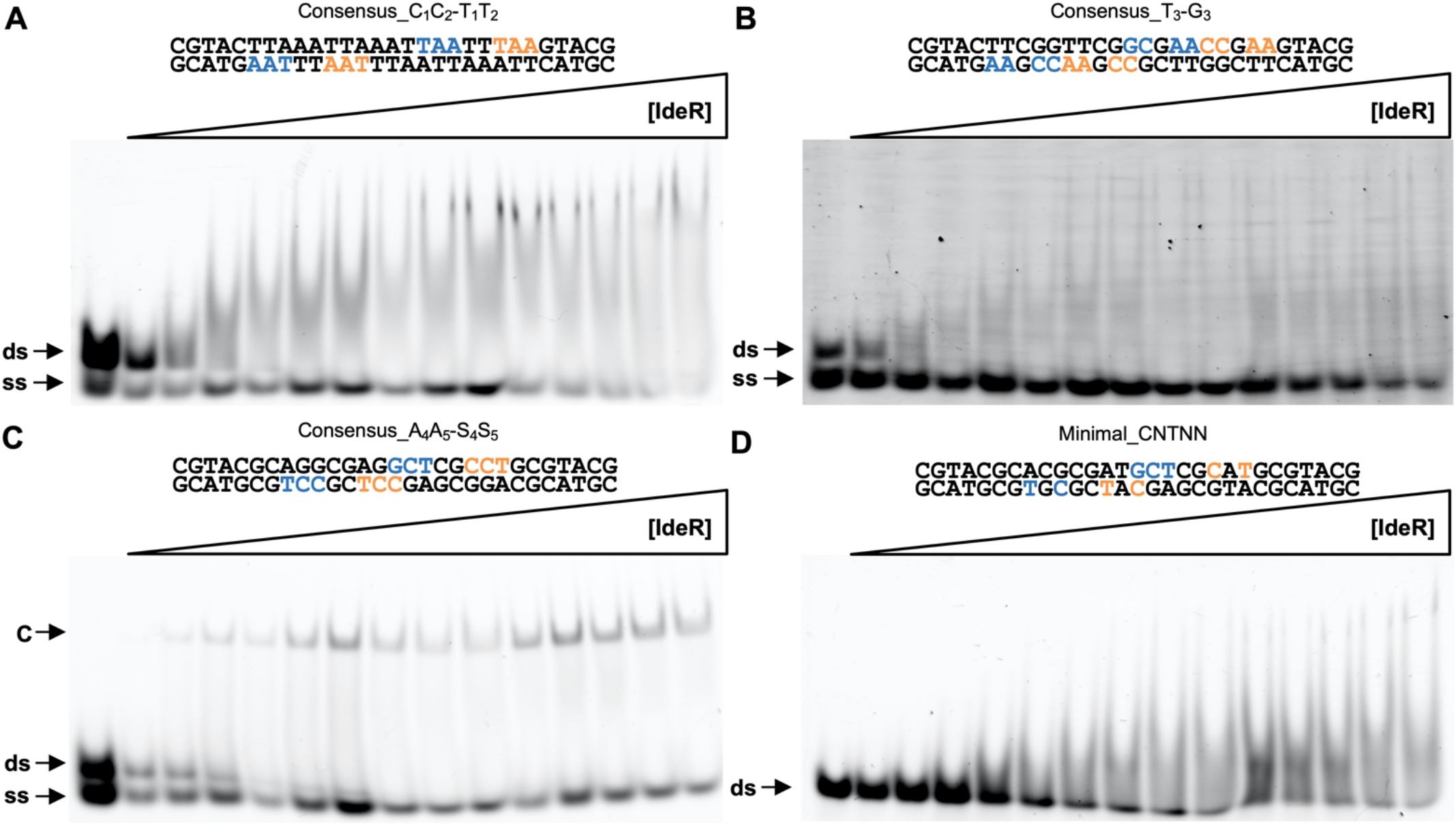
Additional EMSA analysis of IdeR binding to DNA mutants. A Binding of IdeR^WT^ to the Consensus_C_1_C_2_-T_1_T_2_ DNA sequence. B Binding of IdeR^WT^ to the Consensus_T_3_-G_3_ DNA sequence. C Binding of IdeR^WT^ to the Consensus_A_4_A_5_-S_4_S_5_ DNA sequence. D Binding of IdeR^WT^ to the Minimal_CNTNN DNA sequence. IdeR was added in increasing concentrations (panels A, B and D: 150 nM - 22.5 μM dimer; panel C: 15 nM - 2.25 μM dimer) to 30 nM fluorescence-labeled double-stranded DNA probe in the presence of 30 μM Co^2+^ and competitor DNA. Note that IdeR concentrations used in panels A, B and D are 10-fold higher compared to Fig 4D-F. Protein-DNA complexes were resolved on a 4% Tris-acetate polyacrylamide gel. The left-most lane is a control reaction without protein. ds, unbound double-stranded DNA probe; ss, non-hybridized single-stranded DNA; C, protein-dsDNA complex.

**Figure EV5.**
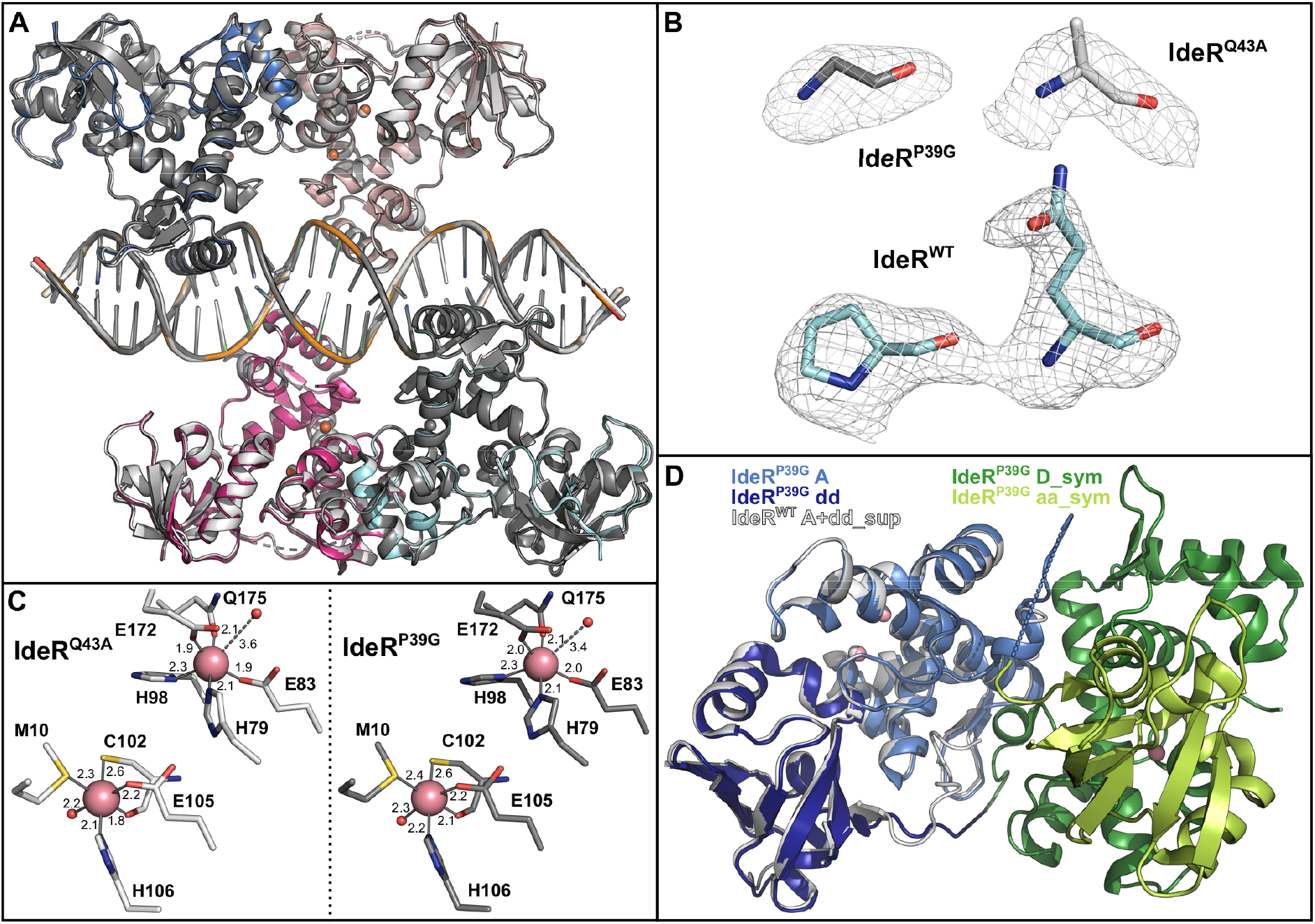
Crystal structures of IdeR variants in complex with the consensus DNA-binding sequence. A Superposition of the Co^2+^-activated complexes of IdeR^Q43A^ (light grey) and IdeR^P39G^ (dark grey) and the Fe^2+^-activated IdeR^WT^ complex (colored by chain) with the consensus DNA-binding sequence. B *mF_o_-DF_c_* omit electron density for G39 in subunit D of IdeR^P39G^ (dark grey) and A43 in subunit A of IdeR^Q43A^ (light grey), compared to *mF_o_-DF_c_* omit electron density for P39 and Q43 in subunit B of the Fe^2+^-activated IdeR^WT^-consensus DNA complex (cyan), shown as grey mesh contoured at +3 σ. C The metal-binding sites in subunit B of Co^2+^-activated IdeR^Q43A^ (light grey, left panel) and subunit B of IdeR^P39G^ (dark grey, right panel). Oxidation of the primary site ligand Cys102 has not occurred to a significant degree in either crystal. Metal-ligand bonds are indicated by grey lines, the dashed line between the ancillary site metal ion and water ligand indicating a long, weak bond. Bond distances are given in Å. D The SH3-like domain swap takes place in IdeR^P39G^, but the loop connecting the dimerization domain and the swapped SH3-like domain assumes a different conformation than in the other IdeR-DNA complexes. Shown are IdeR^P39G^ chain A (marine), consisting of the DNA-binding and dimerization domains, and the associated swapped SH3-like domain (chain dd, dark blue), superimposed with the corresponding IdeR^WT^ chains A and dd (light grey), as well as IdeR^P39G^ chains D_sym and aa_sym (dark green and lime green, respectively) of the symmetry mate which swaps the SH3-like domain with chain A. The connections of the A and aa_sym or D and dd_sym chains are indicated by dashed lines (marine and dark blue, respectively).

