## Appendix Tables S1 and S2 for "The bacterial iron sensor IdeR recognizes its DNA targets by indirect readout"

### Contents

**Appendix Tables**

**Table S1:** Root-mean square distance (RMSD) between aligned Cα atoms of different IdeR structures.

**Table S2:** List of oligonucleotides used in this study.

**Appendix References**

### Appendix Tables

**Table S1. Root-mean square distance (RMSD) between aligned C$\alpha$ atoms of different IdeR structures.** Structures were superimposed using the Coot (Emsley *et al*, 2010) secondary structure matching (SSM) algorithm. RMSD values are given in Å and the number of aligned residues is given in parentheses. IdeR$^{WT}$, IdeR$^{Q43A}$ and IdeR$^{P39G}$ refers to IdeR from *Saccharopolyspora erythraea* and engineered variants thereof (structures determined in this work). consDNA, consensus DNA; C10S1DNA, cluster 10 site 1 DNA; *Mt*IdeR, *Mycobacterium tuberculosis* IdeR in complex with $Co^{2+}$ and DNA (PDB ID 1U8R) (Wisedchaisri *et al*, 2004); *Cd*DtxR, *Corynebacterium diphtheriae* DtxR in complex with $Co^{2+}$ and DNA (PDB ID 1C0W) (Pohl *et al*, 1999).

| | **$Co^{2+}$-IdeR$^{WT}$** | **$Co^{2+}$-IdeR$^{WT}$ + consDNA** | **$Fe^{2+}$-IdeR$^{WT}$ + consDNA** | **$Co^{2+}$-IdeR$^{WT}$ + C10S1DNA** | **$Co^{2+}$-IdeR$^{Q43A}$ + consDNA** | **$Co^{2+}$-IdeR$^{P39G}$ + consDNA** | **$Co^{2+}$-*Mt*IdeR + DNA** | **$Co^{2+}$-*Cd*DtxR + DNA** |
|---|---|---|---|---|---|---|---|---|
| **$Co^{2+}$-IdeR$^{WT}$** | - | 0.79 (434) | 0.75 (436) | 0.80 (432) | 0.80 (436) | 0.76 (432) | 1.28 (421) | 1.53 (373) |
| **$Co^{2+}$-IdeR$^{WT}$ + consDNA** | 0.79 (434) | - | 0.35 (909) | 0.35 (902) | 0.43 (906) | 0.59 (889) | 1.21 (851) | 1.26 (715) |
| **$Fe^{2+}$-IdeR$^{WT}$ + consDNA** | 0.75 (436) | 0.35 (909) | - | 0.30 (902) | 0.28 (914) | 0.52 (891) | 1.18 (860) | 1.25 (713) |
| **$Co^{2+}$-IdeR$^{WT}$ + C10S1DNA** | 0.80 (432) | 0.35 (902) | 0.30 (902) | - | 0.32 (902) | 0.58 (884) | 1.20 (855) | 1.28 (717) |
| **$Co^{2+}$-IdeR$^{Q43A}$ + consDNA** | 0.80 (436) | 0.43 (906) | 0.28 (914) | 0.33 (902) | - | 0.66 (889) | 1.20 (859) | 1.29 (720) |
| **$Co^{2+}$-IdeR$^{P39G}$ + consDNA** | 0.76 (432) | 0.59 (889) | 0.52 (891) | 0.58 (884) | 0.66 (889) | - | 1.14 (852) | 1.41 (714) |
| **$Co^{2+}$-*Mt*IdeR + DNA** | 1.28 (421) | 1.21 (851) | 1.17 (860) | 1.20 (855) | 1.20 (859) | 1.14 (852) | - | 1.35 (720) |
| **$Co^{2+}$-*Cd*DtxR + DNA** | 1.53 (373) | 1.26 (715) | 1.25 (713) | 1.28 (717) | 1.29 (720) | 1.41 (714) | 1.35 (720) | - |

**Table S2. List of oligonucleotides used in this study.**

| Name | Sequence 5'→3' | Description |
|---|---|---|
| IdeR_NdeI_fwd | TATCATATGAACGATCTCATCGATACCACCGAGATG | Forward primer for cloning the *S. erythraea ideR* gene into pET-28a-TEV (*Nde*I restriction site underlined) |
| IdeR_Hind_stop_ rev | TATAAGCTTTCACTTGACGCGCACCATGACCG | Reverse primer for cloning the *S. erythraea ideR* gene into pET-28a-TEV (*Hind*III restriction site underlined) |
| IdeR_Q43A_fwd | GAGCGGCCCCACGGTGAGCGCGACGGTC | Forward mutagenic primer to create the Q43A mutation in *S. erythraea ideR* inserted into pET-28a-TEV (mutated nucleotides underlined) |
| IdeR_Q43A_rev | CTCCATCCGCGCGACCGTCGCGCTCACC | Reverse mutagenic primer to create the Q43A mutation in *S. erythraea ideR* inserted into pET-28a-TEV (mutated nucleotides underlined) |
| IdeR_P39G_fwd | GGAGCAGAGCGGCGGCACGGTGAGCCAG | Forward mutagenic primer to create the P39G mutation in *S. erythraea ideR* inserted into pET-28a-TEV (mutated nucleotides underlined) |
| IdeR_P39G_rev | CTGGCTCACCGTGCCGCCGCTCTGCTCC | Reverse mutagenic primer to create the P39G mutation in *S. erythraea ideR* inserted into pET-28a-TEV (mutated nucleotides underlined) |
| C10S1_FAM_fwd | [FAM]-CGTACTTTGGTAAAGCTAACCTAAGTCACC | Forward strand of the cluster 10 site 1 sequence used for EMSA analyses (5' FAM label) |
| C10S1_rev | GGTGACTTAGGTTAGCTTTACCAAAGTACG | Reverse strand of the cluster 10 site 1 sequence used for EMSA analyses |
| C10S2_Cy5_fwd | [Cy5]-GATGCTTAGGTTAGCCTACCCGACGCTGA | Forward strand of the cluster 10 site 2 sequence used for EMSA analyses (5' Cy5 label) |
| C10S2_rev | TCAGCGTCGGGTAGGCTAACCTAAGCATC | Reverse strand of the cluster 10 site 2 sequence used for EMSA analyses |
| C23S1_Cy5_fwd | [Cy5]-AGTCATAAAGTTAGGTTCGCCTCACTACT | Forward strand of the cluster 23 sequence used for EMSA analyses (5' Cy5 label) |
| C23S1_rev | AGTAGTGAGGCGAACCTAACTTTATGACT | Reverse strand of the cluster 23 sequence used for EMSA analyses |
| Half_Cy5_fwd | [Cy5]-CGTACCCGGTTTAGGGCCACCTAAGTACG | Forward strand of the half site sequence used for EMSA analyses (5' Cy5 label) |
| Half_rev | CGTACTTAGGTGGCCCTAAACCGGGTACG | Reverse strand of the half site sequence used for EMSA analyses |
| IdeR_Cons_Cy5_ fwd | [Cy5]-CGTACTTAGGTTAGGCTAACCTAAGTACG | Forward strand of the Consensus_Full sequence used for EMSA analyses (5' Cy5 label) |
| IdeR_Cons_FAM_fwd | [FAM]-CGTACTTAGGTTAGGCTAACCTAAGTACG | Forward strand of the Consensus_Full sequence used for EMSA analyses (5' FAM label) |
| IdeR_Cons_rev | CGTACTTAGGTTAGCCTAACCTAAGTACG | Reverse strand of the Consensus_Full sequence used for EMSA analyses |
| C1-T1_Cy5_fwd | [Cy5]-CGTACTTAGATTAGACTAATCTAAGTACG | Forward strand of the Consensus_$C_1$-$T_1$ sequence used for EMSA analyses (5' Cy5 label) |
| C1-T1_rev | CGTACTTAGATTAGTCTAATCTAAGTACG | Reverse strand of the Consensus_$C_1$-$T_1$ sequence used for EMSA analyses |
| C2-T2_FAM_fwd | [FAM]-CGTACTTAAGTTAAGTTAACTTAAGTCACG | Forward strand of the Consensus_$C_2$-$T_2$ sequence used for EMSA analyses (5' FAM label) |
| C2-T2_rev | CGTGACTTAAGTTAACTTAACTTAAGTACG | Reverse strand of the Consensus_C2-T2 sequence used for EMSA analyses |

**Table S2 continued.**

| Name | Sequence 5'→3' | Description |
|---|---|---|
| C1C2-T1T2_Cy5_fwd | [Cy5]-CGTACTTAAATTAAATTAATTTAAGTACG | Forward strand of the Consensus_C1C2-T1T2 sequence used for EMSA analyses (5' Cy5 label) |
| C1C2-T1T2_rev | CGTACTTAAATTAATTTAATTTAAGTACG | Reverse strand of the Consensus_C1C2-T1T2 sequence used for EMSA analyses |
| T3-G3_Cy5_fwd | [Cy5]-CGTACTTCGGTTCGGCGAACCGAAGTACG | Forward strand of the Consensus_T3-G3 sequence used for EMSA analyses (5' Cy5 label) |
| T3-G3_rev | CGTACTTCGGTTCGCCGAACCGAAGTACG | Reverse strand of the Consensus_T3-G3 sequence used for EMSA analyses |
| A4A5-S4S5_Cy5_fwd | [Cy5]-CGTACGCAGGCGAGGCTCGCCTGCGTACG | Forward strand of the Consensus_$A_4A_5$-$S_4S_5$ sequence used for EMSA analyses (5' Cy5 label) |
| A4A5-S4S5_rev | CGTACGCAGGCGAGCCTCGCCTGCGTACG | Reverse strand of the Consensus_$A_4A_5$-$S_4S_5$ sequence used for EMSA analyses |
| CNTNN_Cy5_fwd | [Cy5]-CGTACGCACGCGATGCTCGCATCCGTACG | Forward strand of the Minimal_CNTNN sequence used for EMSA analyses (5' Cy5 label) |
| CNTNN_rev | CGTACGGATGCGAGCATCGCGTGCGTACG | Reverse strand of the Minimal_CNTNN sequence used for EMSA analyses |
| Cons_long_fwd | CGTACTTAGGTTAGGCTAACCTAAGTCACG | Forward strand of the Consensus_Full sequence used for co-crystallization with $IdeR^{WT}$ and $IdeR^{P39G}$ |
| Cons_long_rev | CGTGACTTAGGTTAGCCTAACCTAAGTACG | Reverse strand of the Consensus_Full sequence used for co-crystallization with $IdeR^{WT}$ and $IdeR^{P39G}$ |
| Cons_short_fwd | CGTACTTAGGTTAGGCTAACCTAAGTACG | Forward strand of the Consensus_Full sequence used for co-crystallization with $IdeR^{Q43A}$ |
| Cons_short_rev | CGTACTTAGGTTAGCCTAACCTAAGTACG | Reverse strand of the Consensus_Full sequence used for co-crystallization with $IdeR^{Q43A}$ |
| C10S1_xtal_fwd | CGTACTTTGGTAAAGCTAACCTAAGTCACC | Forward strand of the Cluster 10 S1 sequence used for co-crystallization with $IdeR^{WT}$ |
| C10S1_xtal_rev | GGTGACTTAGGTTAGCTTTACCAAAGTACG | Reverse strand of the Cluster 10 S1 sequence used for co-crystallization with $IdeR^{WT}$ |
