## Supplementary material for "The bacterial iron sensor IdeR recognizes its DNA targets by indirect readout": Table EV1

**Table EV1. Clusters potentially regulated by IdeR in the genome of *S. erythraea* and putative IdeR binding sites and relevant genes found in each cluster.**

[a]Closest potentially regulated gene(s). [b]Position with respect to annotated start codon.

| Cluster number | Cluster span | Binding site | Sequence NNNNNTTAGGTTAGSCTAACCTAANNNNN | Closest gene(s)[a] | Position[b] | Relevant or metal-related genes in the cluster |
|---|---|---|---|---|---|---|
| 1 | SACE_0130-SACE_0134 | C01S1 | AAGCCTGAGGTTTGCCTAACCGAAATAAG | SACE_0130 | -88 | SACE_0130: Bacterioferritin B |
| | | C01S2 | CGAAATAAGGTGAACAAAACCTATATAAG | SACE_0130 | -68 | |
| | | C01S3 | GTCATCTAAGTTAGGGTTACCTAAGTTGG | SACE_0134 | -9 | |
| 2 | SACE_0257-SACE_0262 | C02S1 | TCCCCTTAGGTCGCCCTAAGTAGCTGGAT | SACE_0257 | -155 | SACE_0257: Peptide/Nickel ABC transport system, substrate-binding component |
| | | | | | | SACE_0258: Peptide/Nickel ABC transport system, permease component |
| | | | | | | SACE_0259: Peptide/Nickel ABC transport system, permease component |
| | | | | | | SACE_0260: Peptide/Nickel ABC transport system, ATP-binding component |
| | | C02S2 | AAATGCCAGGTTAGTCTCACGTCACTCTG | SACE_0257 | -99 | SACE_0261: Peptide/Nickel ABC transport system, ATP-binding component |
| 3 | SACE_0411-SACE_0415 | C03S1 | TAGTCTTAGGCCGACCTAACCTAGGTGCG | SACE_0411 | -41 | SACE_0411: fepD, ferrichrome ABC transport system, permease component |
| | | | | | | SACE_0412: fepG, $Fe^{3+}$ enterobactin ABC transport system, permease component |
| | | | | | | SACE_0413: fepC, $Fe^{3+}$ enterobactin ABC transport system, ATP-binding component |
| 4 | SACE_0613-SACE_0618 | C04S1 | GGGTATTAGGTTAGCCTAACCTGCGAGCA | SACE_0613 | -172 | SACE_0617: DsbA-like thioredoxin domain protein |
| | | | | SACE_0614 | -221 | SACE_0618: Integral membrane cytochrome biogenesis protein |
| | | | | | | SACE_0619: sodF, superoxide dismutase [Fe-Zn] I, Fe-Mn family |
| | | | | | | SACE_0620: Iron complex ABC transport system, substrate-binding component |
| | | | | | | SACE_0621:Iron complex ABC transport system, permease component |
| | | | | | | SACE_0622: Iron complex ABC transport system, ATP-binding component |
| 5 | SACE_1004-SACE_1011 | C05S1 | ACGGGTTAGGTTCGCCTAGCCTGTCCGCG | SACE_1004 | -33 | SACE_1010: 4Fe-4S ferredoxin |
| | | | | SACE_1005 | -254 | |

| Cluster number | Cluster span | Binding site | Sequence NNNNNTTAGGTTAGSCTAACCTAANNNNN | Closest gene(s)[a] | Position[b] | Relevant or metal-related genes in the cluster |
|---|---|---|---|---|---|---|
| **6** | SACE_1287-SACE_1312 | C06S1 | AAGCATTAGGTTAGGCTCCCCAAACCAGT | SACE_1287 | -26 | SACE_1287: Iron complex ABC transport system, substrate-binding component |
| | | C06S2 | CACCATCCGGTGAGGACCACCTAACCCAG | SACE_1295 | -17 | SACE_1305: Putative NRPS |
| | | C06S3 | AACAATTAGGCGAGGCTAACGTGCGTCGA | SACE_1301 | -13 | SACE_1308: Siderophore-interacting protein |
| | | | | SACE_1300 | -171 | |
| | | C06S4 | AAGTGTTAGGTTAGCCTCAGCTCAGCGGG | SACE_1304 | -22 | |
| | | | | SACE_1303 | -234 | |
| | | C06S5 | TGAACCAAGGTTAGCCTTACTTTTATTGG | SACE_1305 | -266 | |
| | | C06S6 | CTATCTATGGTTAGGCTAGCCTTAATCAA | SACE_1307 | -36 | |
| **7** | SACE_1670-SACE_1676 | C07S1 | GTTGTTTAGGCGAGCCTTCCCTTGCTGGC | SACE_1675 | -163 | SACE_1672: NitT/TauT family ABC transport system, permease component |
| | | | | SACE_1674 | 31 | SACE_1673: NitT/TauT family ABC transport system, substrate-binding component |
| | | | | | | SACE_1674: NitT/TauT family ABC transport system, ATP-binding component |
| **8** | SACE_2001-SACE_2007 | C08S1 | GGATCGCTGGTAAGGCATGCCTAACATGG | SACE_2001 | -94 | SACE_2001: Ferredoxin/flavodoxin $NADP^+$ reductase |
| | | | | SACE_2002 | -207 | |
| **9** | SACE_2401-SACE_2414 | C09S1 | GAGAGTAAGGTGAGGCTTACCTAGCATTT | SACE_2412 | -31 | SACE_2412: High-affinity iron transporter |
| | | | | SACE_2411 | -159 | SACE_2413: efeO, iron uptake system component |
| **10** | SACE_2687-SACE_2704 | C10S1 | GTGACTTAGGTTAGCTTTACCAAAGTACG | SACE_2689 | -103 | SACE_2689: trpE, anthranilate synthase component I |
| | | | | SACE_2690 | -288 | SACE_2690: Enterobactin MFS exporter, ENTS family |
| | | | | | | SACE_2692: MbtH-like protein |
| | | | | | | SACE_2693: NRPS, mycobactin phenyloxazoline synthetase |
| | | | | | | SACE_2694: Mycobactin salicyl-AMP ligase |
| | | | | | | SACE_2695: Putative NRPS |

| Cluster number | Cluster span | Binding site | Sequence NNNNNTTAGGTTAGSCTAACCTAANNNNN | Closest gene(s)[a] | Position[b] | Relevant or metal-related genes in the cluster |
|---|---|---|---|---|---|---|
| | | C10S2 | GATGCTTAGGTTAGCCTACCCGACGCTGA | SACE_2690 | -123 | SACE_2696: Putative NRPS |
| | | | | SACE_2689 | -268 | SACE_2697: Putative $Fe^{3+}$-siderophore ABC transport system, substrate-binding component |
| | | C10S3 | GCAACATAGCATAGCCTCACCTAACTAGC | SACE_2698 | -18 | |
| | | | | SACE_2697 | -270 | |
| **11** | SACE_2917-SACE_2922 | C11S1 | TCAGTTTAGCTTAGGTTAACCTAACCGCC | SACE_2921 | -34 | SACE_2921: Iron complex ABC transport system, substrate-binding component |
| | | | | SACE_2922 | -515 | |
| **12** | SACE_3033-SACE_3042 | C12S1 | ACGGGCTAAGTTAGTCATGCCTAAATAAA | SACE_3033 | -136 | SACE_3034: Iron complex ABC transport system, substrate-binding component |
| | | | | | | SACE_3035: Putative NRPS |
| | | | | | | SACE_3036: MbtH protein |
| | | C12S2 | CTAAATAAAGGTAGGCTCACCTTCACCGC | SACE_3033 | -116 | SACE_3037: Putative ABC transport system, ATP-binding component |
| | | C12S3 | ATCTTTTAGAGTAGGCTTACCTAACACTT | SACE_3038 | -117 | SACE_3038: Putative ABC transport system, ATP-binding component |
| | | | | SACE_3039 | -145 | |
| **13** | SACE_3145-SACE_3150 | C13S1 | CATCCGCGGGTTACCGCGACGTAACCCTC | SACE_3146 | -153 | SACE_3145: [illegible], ferredoxin reductase, 3-phenylpropionate/trans-cinnamate dioxygenase ferredoxin component |
| | | | | SACE_3147 | -149 | SACE_3146: hcaC2, 3-phenylpropionate/trans-cinnamate dioxygenase ferredoxin component |
| | | C13S2 | ACTTTGCAGGTTAGGGCCACCTTCCGGCC | SACE_3147 | -41 | SACE_3150: Ferredoxin subunit of nitrite reductase and ring-hydroxylating dioxygenase |
| | | | | SACE_3146 | -261 | |
| **14** | SACE_3570-SACE_3575 | C14S1 | TGAGGGTTGGTTAGGCTAGGCTACCTTTT | SACE_3571 | -83 | SACE_3574: UDP-N-acetylmuramoylalanine--D-glutamate ligase |
| | | | | SACE_3570 | -128 | SACE_3575: cobQ-2, cobyric acid synthase |
| | | C14S2 | CTTGCTTTCGGCGGGCTAATACGCGCCTG | SACE_3574 | -122 | |
| | | | | SACE_3573 | -307 | |
| **15** | SACE_3852-SACE_3855 | C15S1 | CTGACCAAGGTAAGGCTAGCCTGAGTAAA | SACE_3855 | -14 | SACE_3852: dhbB, isochorismatase (2,3 dihydro-2,3 dihydroxybenzoate synthase) |

| Cluster number | Cluster span | Binding site | Sequence NNNNNTTAGGTTAGSCTAACCTAANNNNN | Closest gene(s)[a] | Position[b] | Relevant or metal-related genes in the cluster |
|---|---|---|---|---|---|---|
| | | | | | | SACE_3853: dhbE, 2,3-dihydroxybenzoate-AMP ligase |
| | | | | | | SACE_3854: dhbC, isochorismate synthase |
| | | | | | | SACE_3855: dhbA, 2,3-dihydro-2,3-dihydroxybenzoate dehydrogenase |
| **16** | SACE_4076-SACE_4077 | C16S1 | GCGTGCTAGCTTAGCCTAACCTTAATGCT | SACE_4076 | -13 | SACE_4076: sidF, ferrichrome ABC transport system, substrate-binding protein |
| | | | | SACE_4077 | -35 | SACE_4077: sidE, siderophore-interacting protein |
| **17** | SACE_4272_SACE_4282 | C17S1 | CGCCGCAAAAGCAGGCTTGCCTAAGTTCG | SACE_4278 | -59 | SACE_4273: Spectinomycin phosphotransferase |
| | | | | SACE_4277 | -133 | SACE_4275: Thioesterase involved in non-ribosomal peptide biosynthesis |
| | | | | | | SACE_4276: Oleandomycin ABC transport system, permease component |
| | | | | | | SACE_4277: Oleandomycin ABC transport system, ATP-binding component |
| | | | | | | SACE_4279: NRPS/PKS |
| **18** | SACE_4972-SACE_4974 | C18S1 | ATAATTAAGGTTTGCCTTTCCTTGGTGCT | SACE_4973 | -69 | SACE_4972: Cation-binding protein, hemerythrin HHE family |
| | | | | SACE_4972 | -112 | SACE_4973: Bacterioferritin B |
| **19** | SACE_5040-SACE_5045 | C19S1 | CAGGCACAGCTGAGCCTGCCCTACGCGGT | SACE_5044 | -441 | SACE_5042-SACE_5044: $Fe^{3+}$ ABC transport system |
| | | | | SACE_5045 | 972 | |
| **20** | SACE_5662-SACE_5667 | C20S1 | ACTACTTAGGTTAGCCTTCCTTATATTCG | SACE_5666 | -41 | SACE_5666: Iron complex ABC transport system, substrate-binding component |
| | | | | SACE_5665 | -180 | |
| **21** | SACE_5741-SACE_5745 | C21S1 | CCGGCGACTCCTCGGTTAACCGAAGGATG | SACE_5742 | -219 | SACE_5743: Iron complex ABC transport system, substrate-binding component |
| | | | | SACE_5743 | -332 | SACE_5744: Iron complex ABC transport system, permease component |
| | | | | | | SACE_5745: Iron complex ABC transport system, ATP-binding component |
| **22** | SACE_5876-SACE_5887 | C22S1 | ATCATGCAGGTTAGGCTTGCCTAACTATG | SACE_5883 | 3 | SACE_5882: Ribosomal protein S12 methylthiotransferase |
| | | | | SACE_5882 | -218 | SACE_5883: Amino-acid acetyltransferase |

| Cluster number | Cluster span | Binding site | Sequence NNNNNTTAGGTTAGSCTAACCTAANNNNN | Closest gene(s)[a] | Position[b] | Relevant or metal-related genes in the cluster |
|---|---|---|---|---|---|---|
| | | | | | | SACE_5884: Amino-acid N-acetyltransferase |
| | | | | | | SACE_5887: lldD2, L-lactate dehydrogenase (cytochrome) |
| **23** | SACE_6887-SACE_6902 | C23S1 | AGTCATAAAGTTAGGTTCGCCTCACTACT | SACE_6902 | -90 | SACE_6889: nuoN, NADH dehydrogenase (quinone), subunit N |
| | | | | | | SACE_6890: nuoM, NADH dehydrogenase (quinone), subunit M |
| | | | | | | SACE_6891: nuoL, NADH dehydrogenase (quinone), subunit L |
| | | | | | | SACE_6892: nuoK, NADH dehydrogenase (quinone), subunit K |
| | | | | | | SACE_6893: nuoJ, NADH dehydrogenase (quinone), subunit J |
| | | | | | | SACE_6894: nuoI, NADH dehydrogenase (quinone), subunit I |
| | | | | | | SACE_6895: nuoH, NADH dehydrogenase (quinone), subunit H |
| | | | | | | SACE_6896: nuoG, NADH dehydrogenase (quinone), subunit G |
| | | | | | | SACE_6897: nuoF, NADH dehydrogenase (quinone), subunit F |
| | | | | | | SACE_6898: nuoE, NADH dehydrogenase (quinone), subunit E |
| | | | | | | SACE_6899: nuoD, NADH dehydrogenase (quinone), subunit D |
| | | | | | | SACE_6900: nuoC, NADH dehydrogenase (quinone), subunit C |
| | | | | | | SACE_6901: nuoB, NADH dehydrogenase (quinone), subunit B |
| | | | | | | SACE_6902: nuoA, NADH dehydrogenase (quinone), subunit A |
